# Genomic analysis of *Ostreococcus tauri*-infecting viruses reveals a hypervariable region associated with host–virus interactions

**DOI:** 10.1101/2025.08.15.670463

**Authors:** Julie Thomy, Julien Henri, David Demory, Frederic Sanchez, Marie-Line Escande, Nigel Grimsley, Sheree Yau

## Abstract

While the genus *Prasinovirus*, known to infect prasinophyte green algae (class Mamiellophyceae), is abundant in the oceans, the genetic mechanisms governing the ecology and (co)evolution of these viruses remain poorly understood. In this study, we sequenced the complete genomes of eighteen viruses infecting the cosmopolitan unicellular green alga *Ostreococcus tauri* (OtVs) and one specific to *Micromonas commoda* (McV-20T). Of these viruses, twelve were previously isolated from the coastal Mediterranean Sea and seven were newly isolated from two different geographical locations (i.e., the South Pacific Gyre and the North Sea). Phylogenetic analysis classified these viruses as new members of the *Prasinovirus* genus and defined three distinct OtV clades: designated OtV-type 1, OtV-type 2a and OtV-type 2b. The OtV-type 1 includes three new viruses isolated from the Pacific Ocean and the previously sequenced genome OtV6. The OtV-type 1 form a large cluster within the *Micromonas*-infecting virus clade, sharing five unique homologous genes with all *Micromonas* viruses. In addition, genetic features of OtV-type 1, including a higher number of CDSs (∼260) and lower GC content (∼41%), were more closely allied to *Micromonas* viruses than to those of OtVs, suggesting an alternative host or a recent host switch from *Micromonas* to *O. tauri.* By analyzing the OtV genomes, we found a faster-evolving central hypervariable region (HVR), where the OtV-type 2b displayed the largest region, i.e., three times longer than other OtVs. This region encodes genes mainly associated with host cell recognition and attachment and under strong selective pressure (positive and negative). Notably, most viruses associated with OtV-type 2b showed the broadest host range. Infection dynamics between the hosts and the viruses appeared highly specific to host–virus pairs, suggesting complex interactions in the *Ostreococcus*-prasinovirus system. Finally, by observing viral lysates with electron microscopy, we observed novel morphologies never described for these viruses. Overall, this study provides new insights into the genetic diversity of prasinoviruses and describes for the first time a viral plasticity that might be strongly shaped by antagonistic coevolution with their hosts.

## Introduction

Viruses are described as the most common and diverse parasites in the oceans (Suttle 2005, 2007). Through cell lysis, marine viruses play a major role in marine ecosystems and microbial food webs by facilitating recycling of organic matter and nutrients (Fuhrman 1999; Wilhelm & Suttle 1999; Guidi et al. 2016; Middelboe & Brussaard 2017; Mayers et al. 2023). Beyond mortality, they also strongly shape the evolution of their host (Thingstad 2000; Weinbauer 2004; Friman & Buckling 2013) by driving host microdiversity (i.e., intra genomic diversity) (Fuhrman & Campbell 1998; Moore et al. 1998) through genetic mechanisms, such as mutations, horizontal gene transfers, and viral resistance (Avrani et al. 2011; Marston et al. 2012). The evolution of viral genomes is also strongly shaped by their host (Retel et al. 2019). Viruses and their hosts often undergo antagonistic coevolution, marked by reciprocal genetic exchanges over time (Martiny et al. 2014). These dynamics involve ongoing evolution, where the host evolves mechanisms to resist the antagonist, and the antagonist evolves counter-strategies to overcome the defenses of the host (Van Valen 1974). In this way, the coevolutionary process between viruses and their hosts is also described by complex infection networks (Fortuna et al. 2010; Weitz et al. 2013; Edwards & Hayward 2024). A nested network is one of the hypothesized infection patterns resulting from gene-for-gene coevolution. In this process, viruses evolve to broaden their host range in response to increasing host resistance (Agrawal & Lively 2003; Dennehy 2012; Weitz et al. 2013). These complex interplays have been widely studied in the bacteria– phage model systems (Flores et al. 2011; Stern & Sorek 2011; Weitz et al. 2013) as well as in marine cyanobacteria and cyanophages (Waterbury & Valois 1993; Avrani et al. 2011; Zborowsky & Lindell 2019) and in microbial eukaryote–virus systems (Edwards & Hayward 2024; Frickel et al. 2016, 2018). Phytoplankton have developed various defense mechanisms and strategies to protect themselves against pathogens (Thomas et al. 2012). These strategies are diverse and include life-phase switch mechanisms (Brussaard et al. 2007; M. Frada et al. 2008; M. J. Frada et al. 2017), programmed cell death (Bidle 2016), lipidome plasticity (Schleyer et al. 2023) or spore formation (Pelusi et al. 2021). In the green microalgae *Ostreococcus* spp., antiviral resistance has been associated with changes in the size of small “outlier” chromosomes (SOC), also known as “immune” chromosomes, as well as in the transcription pattern of the latter (Thomas et al. 2011; Derelle et al. 2018; Yau et al. 2016, 2018, 2020). This chromosome encodes many predicted membrane surface proteins, glycoconjugate production proteins important for host–virus interactions, and proteins associated with carbohydrate transport and metabolism, most of them being overexpressed in virus-resistant strains. In addition, these virus-resistant lines showed a reduction in the adsorption of viral particles on the surface of host cells (Yau et al. 2018).

Conversely, another important aspect is to better understand how these coevolutionary dynamics influence the adaptive response of viruses to spontaneous resistance in their hosts. The genus *Prasinovirus* belongs to the phylum *Nucleocytoviricota*, often referred to as “giant viruses”. They infect cosmopolitan unicellular green algae of the class Mamiellophyceae, which includes the genera *Micromonas*, *Ostreococcus*, and *Bathycoccus* (Clerissi et al. 2012; Baudoux et al. 2015; Derelle et al. 2015; Bachy et al. 2021; Listmann et al. 2023), as well as the species *Mantoniella tinhauana* (Rey Redondo et al. 2024). Mamiellophyceae are widespread and dominate the eukaryotic picophytoplankton fraction in coastal waters (Tragin & Vaulot 2018, 2019). Their abundance supports dense populations of prasinoviruses, which are detected throughout marine environments where their hosts have been isolated (M. Cottrell & Suttle 1991; M. T. Cottrell & Suttle 1995; Adriana Zingone et al. 1999; Bellec et al. 2010). Metagenomic analyses further confirm that prasinoviruses are both widespread and abundant in the oceans (Short & Short 2008; Hingamp et al. 2013; Chen et al. 2018; Endo et al. 2020; Farzad et al. 2022; Ha et al. 2023). The global distribution of prasinoviruses and their hosts suggests a long-standing coevolutionary relationship. Indeed, *Prasinovirus*–Mamiellophyceae interactions exhibit a high degree of host specificity, with many viruses infecting only a single species or even specific strains within a species (A. Zingone et al. 2006; Clerissi et al. 2012; Bellec et al. 2014; Baudoux et al. 2015). These interactions are described as nested, wherein specialist viruses (narrow host range) and generalist viruses (broad host range) infect hosts that vary in their susceptibility or resistance to infection. Phylogenetic analyses of prasinoviruses and their hosts often showed congruent evolutionary trees, supporting the notion of virus–host coevolution (Bellec et al. 2014). While a strong co-speciation signal is observed in most *Prasinovirus*–Mamiellophyceae relationships, the *Bathycoccus*–virus system appears to reflect a more complex cophylogenetic history. This includes evidence for an early host- switch from *Bathycoccus* to *Micromonas*, suggesting that prasinovirus colonization of *Micromonas* may be more recent (Bellec et al. 2014). Moreover, recent studies have uncovered unexpected host-switching across diverse algal lineages, challenging existing paradigms of strict host–virus cospeciation (Thomy et al. 2024). These findings highlight the intricate evolutionary dynamics of the *Prasinovirus*– Mamiellophyceae system and underscore the importance of exploring the genetic mechanisms that govern host–virus interactions more broadly.

In this work, we used comparative genomic analysis of eighteen distinct isolated viruses infecting *Ostreococcus tauri* (OtVs) and one infecting *Micromonas commoda*. We combined these results with cross-infections of a panel of distinct Mamiellophyceae strains to unveil the genetic signatures associated with OtV diversification and evolution of host range. Finally, the characterization of the infection dynamics and morphology of these viruses provided further insights into gene-to-phenotype associations.

## Results

### Newly isolated prasinoviruses from distant geographic origins

We sought to expand the diversity of marine viruses in our study group by isolating new viruses from distant sites across the globe. Nine taxonomically diverse eukaryotic phytoplankton strains (Trebouxiophyceae, Mamiellophyceae, Pyramimonadophyceae and Pavlovaceae) were challenged with environmental seawater from two distant coastal sites (South Pacific Ocean Gyre and South in the Eastern North Sea) (Supplementary Table 1). No lytic viruses were detected for Trebouxiophyceae, Pyramimonadophyceae and Pavlovaceae strains in our culture conditions using the volumes of seawater tested. The viral titers in the seawater samples were sufficient to lyse only the Mamiellophyceae strains of the genera, *Micromonas*, *Ostreococcus* and *Bathycoccus*. We successfully isolated viruses on *Ostreococcus tauri* RCC4221 and *Micromonas commoda* RCC827 strains from seawater sampled from three distinct sites in the Pacific Ocean along the Chilean coast and from the North Sea (Supplementary Figure 1). Overall, we isolated six new viruses infecting *Ostreococcus tauri* and one infecting *Micromonas commoda* (Supplementary Figure 1).

PCR amplification of the viral DNA polymerase B gene (*polB*) with degenerate primers and Sanger sequencing supported the taxonomic affiliation of these viruses to the genus *Prasinovirus*. Genomes of these seven new prasinoviruses, along with those of twelve genetically distinct *O. tauri* viruses (OtVs) previously isolated from the coastal Mediterranean Sea (Clerissi et al. 2012), were sequenced to extend our knowledge of prasinovirus diversity.

### Genomic features of prasinoviruses

Nineteen new prasinovirus genomes were successfully assembled from Illumina high-throughput sequences, all of which were found to be linear double-stranded DNA genomes of ∼190 kbp (Supplementary Table 2). From the OtV19-T viral lysates, two separate contigs were successfully recovered with a length of 189 and 200-kbp and with a GC composition of 41.9% and 41.2%, respectively. On each of these contigs, different *polB* genes were present in a single copy, suggesting a co-infection of two distinct viruses (dubbed OtV19-T1 and OtV19-T2) on the same host. The presence of two viral genomes was surprising, as all viruses were isolated by picking off individual virus plaques through three successive rounds of plating. One possibility is that the plaque purification was insufficient to separate them, for example, if the virus particles were closely physically associated such that they were co-purified. Another possibility is that these two genomes might be defective for unknown different but complementary functions and require coinfection to reproduce.

While most viruses described in this study presented a highly similar genome length, comparable with the other *Ostreococcus* virus reference genomes, some of them (4 out of 18 viruses) displayed a significantly different DNA base composition (GC content) and a higher number of coding genes (CDS), separating them into two main groups (Supplementary Figure 2). The first group includes most of the viruses, such as those from the Mediterranean and North Seas. This group encoded ∼247 CDSs with a GC content of 45%. The second group, which included mainly viruses from the Pacific Ocean (four out of five), was distinguished by a higher number of CDSs (∼260) and a lower GC content (∼41%). Its genetic features were more closely allied to those of the newly isolated *Micromonas* virus (McV20-T) (see Supplementary Figure 2). A greater number of tRNA genes were also identified in OtV19-O, OtV19-P, OtV19-T1 (8 tRNAs) as well as in OtV6 (9 tRNAs) compared to other viruses which harbor approximately five tRNAs (Supplementary Table 2). Although five tRNAs (tRNA_Gln_, tRNA_Ile_, tRNA_The_, tRNA_Asn_ and tRNA_Tyr_) were found to be common to all OtVs (except OtV1), two (tRNA_Arg_ and tRNA_Pro_) were unique to these four viruses (OtV19-O, OtV19-P, OtV19-T1 and OtV6). Furthermore, some of these (tRNA_Gln_, tRNA_Tyr_, tRNA_Asn_ and tRNA_Thr_), were found to be clustered together near the 3’ end of the genome. tRNA clusters have already been reported in chlorovirus, as well as in prasinovirus genomes (Moreau et al. 2010; Duncan et al. 2020). Hierarchical clustering based on the presence and absence pattern of tRNA genes showed a close relationship between the three OtV19- O, OtV19-P, OtV19-T1 viruses and OtV6 (Supplementary Figure 3). Likewise, viruses within the same cluster showed synteny of tRNA genes in their genomes. The acquisition and selection of specific tRNAs might enable viruses to adapt to codon usage bias for the synthesis of their own proteins by complementing the pool of tRNAs available from their host cell (Michely et al. 2013; Azam et al. 2024), or it might reflect viral adaptations to host defense mechanisms (Zboroskvy et al., 2025). It is therefore possible that these three new viruses isolated from the Pacific Ocean and OtV6 infect a host with a distinct codon usage bias from *O. tauri* in the environment. Finally, terminal inverted repeats (TIRs) were identified from most OtV viruses (16 out of 18) as well as in the McV-20T (Supplementary Table 2), indicating the successful assembly of these genomes. While the biological role of TIRs during infection remains poorly understood, these features appear to be commonly found in prasinovirus genomes (Derelle et al. 2008; Bachy et al. 2018, 2021; Rey Redondo et al. 2024).

### Distinct evolutionary relationship and gene composition

Phylogenetic reconstruction based on DNA polB and concatenation of nineteen core proteins placed the isolated viruses as new members of *Prasinovirus* (Supplementary Figure 4), which is classified within the order *Algavirales* based on other phylogenetic frameworks (Aylward et al. 2021). The resulting phylogenetic tree grouped the McV20-T within the clade containing *Micromonas* viruses (MpV1, MpVSP1, MpVPL1) (Supplementary Figure 4).

Hierarchical clustering based on the orthogroup pattern (Figure 1) revealed a topology comparable to the multi-protein phylogenetic tree with OtVs separated into two clades (Supplementary Figure 4), with the exception that *Ostreococcus* viruses do not form a single large cluster. In contrast, the orthogroup analysis permitted separation of OtV groups according to other factors shaping the non-core genes or pan-genome, which potentially include specific environmental or host-interacting genes. Hence, based on both genome-wide diversity (Figure 1) and the evolutionary relationships (Supplementary Figure 4), three main OtV groups were defined. The first one dubbed OtV-type 1 includes the three Pacific Ocean viruses (OtV19-O, OtV19-P and OtV19-T1) and OtV6, which form a large cluster within the *Micromonas*-infecting viruses clade. The second group segregated into two distinct sub-groups: OtV-type 2a and OtV-type 2b. OtV-type 2a comprises all OtVs (except OtV2, whose host species was not *Ostreococcus tauri*, but *Ostreococcus* sp. clade B (Weynberg et al. 2011)) and the *Ostreococcus mediterraneus* virus 1 (OmV1) while OtV-type 2b comprises the fast-diverging strains OtV06-12, OtV09-565, OtV09-578, and OtV19-T2 (Figure 1).

**Figure 1.**
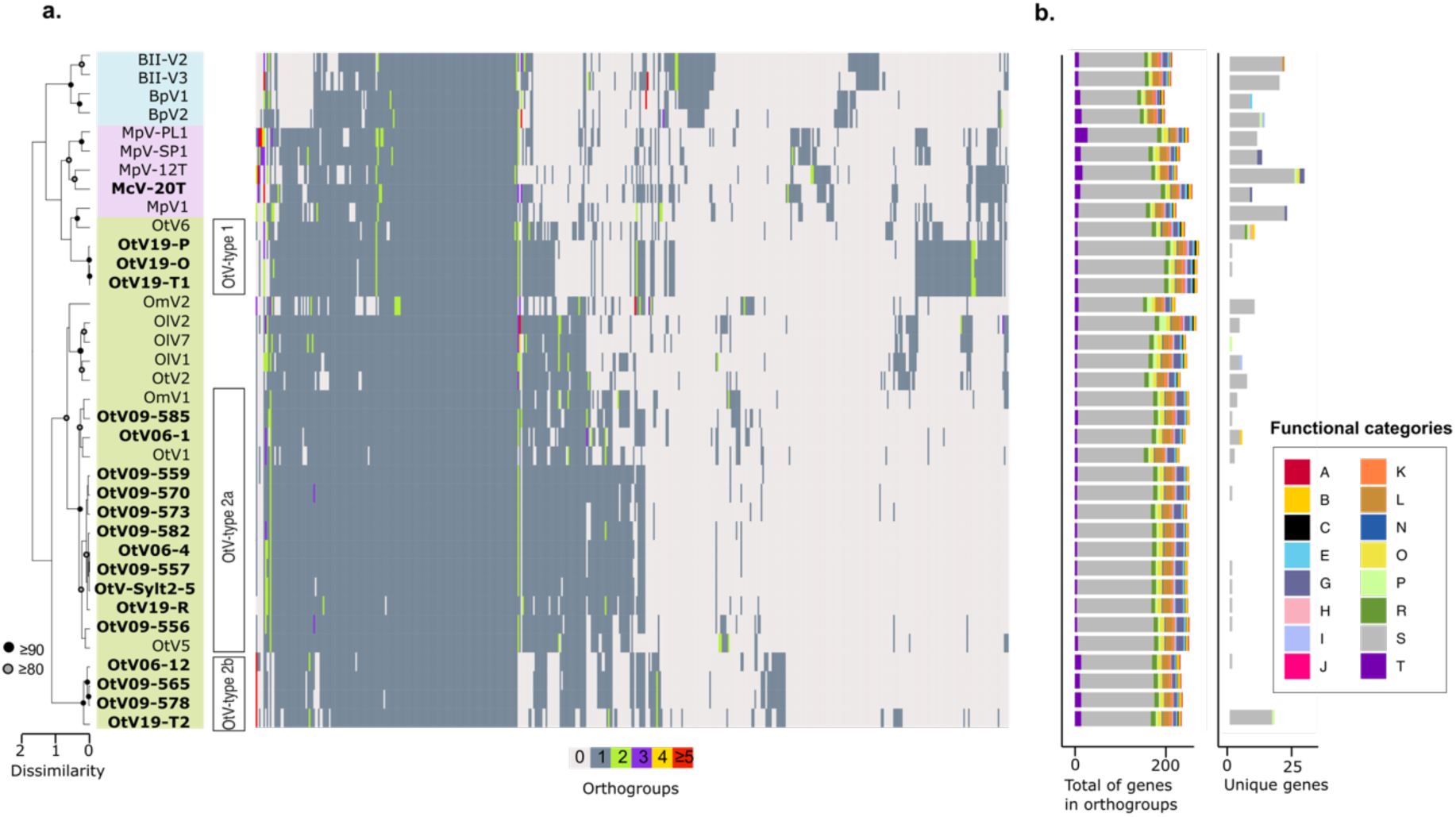
Orthologous protein distribution of prasinoviruses. a. Heatmap showing the distribution and frequency of all orthogroups in *Bathycoccus* (blue background box), *Micromonas* (purple box) and *Ostreococcus* (green box) viruses. Virus names in bold were sequenced in this study. The heatmap was ordered vertically by hierarchical clustering of viral genomes based on the occurrence pattern of all orthogroups (left dendrogram) and horizontally by clustering of the orthogroups based on their distribution in the viral genomes. Nodes marked with a gray dot show genome clusters with over 80% bootstrap support and black dots show over 90% support. b. Bar charts showing (left) the total number of genes assigned to an orthogroup and (right) the number of unique genes (not shared with any other virus analyzed) in each viral genome. Bar charts colored according to the distribution of functional categories based on the EggNOG database.

Pan-genomic analysis revealed that thirty-seven of the 482 orthogroups were present in all prasinovirus genomes (core orthogroups) (Supplementary Table 3). The large *Ostreococcus* virus clade shared a total of 118 core orthogroups, while 136 were found in *Micromonas* virus and 134 in *Bathycoccus* virus genomes. 139 core orthogroups were found among the three newly defined OtV groups. Based on the EggNOG classification, only ∼28% of the genes were assigned to putative functions. Most of them were associated with RNA processing and modification, amino acid metabolism and transport, nucleotide metabolism and transport, lipid metabolism, and DNA replication/repair (Figure 1). Among the *O. tauri* virus clade, OtV-type 1 was the most distinct group with forty-three unique orthogroups and twenty-five of which were present in at least one *Micromonas* virus genome, representing approximately 60% of the orthogroups in this group (Supplementary Table 3). Of all these genes, only six encode known functions, such as a mannitol dehydrogenase C-terminal domain protein, a glycerophosphoryl diester phosphodiesterase family protein, a mitochondrial 18 kDa protein and an alternative oxidase (AOX), a copper/zinc superoxide dismutase, and an endosialidase chaperone. Finally, OtV-type 2a and OtV-type 2b share six and eleven unique orthogroups, respectively. Interestingly, within OtV-type 2b, two signature genes were annotated as glycosyltransferase family 17 and a coiled stalk of trimeric autotransporter adhesin, both involved in host interaction and immune evasion.

### The OtV genome is modular comprising syntenic and variable regions

Overall, the genomes of OtVs displayed a high collinearity and gene synteny (Figure 2). The pairwise average nucleotide identity (ANI) values between them ranged from approximately 71% to 99%, indicating a very low level of divergence (Supplementary Table 4). The organization and structure of gene function followed a genomic pattern of locally collinear blocks (LCBs) (Darling et al. 2004), which define homologous regions of sequence shared by the genomes (Figure 2). The two first LCBs (LCB1 and LCB2) included a broad range of genes associated with carbohydrate, nucleotide and amino acid biosynthesis and metabolism. The LCB3 and LCB4 included genes with functions involved in adhesion, cell recognition and morphogenesis. The central LCBs (5 to 9) were very conserved among the prasinoviruses and contained the bulk of the core genes associated with DNA transcription, replication and post transcriptional modification. The LCB10 and LCB11 encoded genes involved in amino acid and polysaccharide synthesis and metabolism, while the last four blocks also contained genes associated to the DNA replication, morphogenesis including two capsid genes and DNA polymerase B and DNA topoisomerase II. In addition, some glycosyltransferases, associated with carbohydrate biosynthesis and metabolism, were found in these regions.

**Figure 2.**
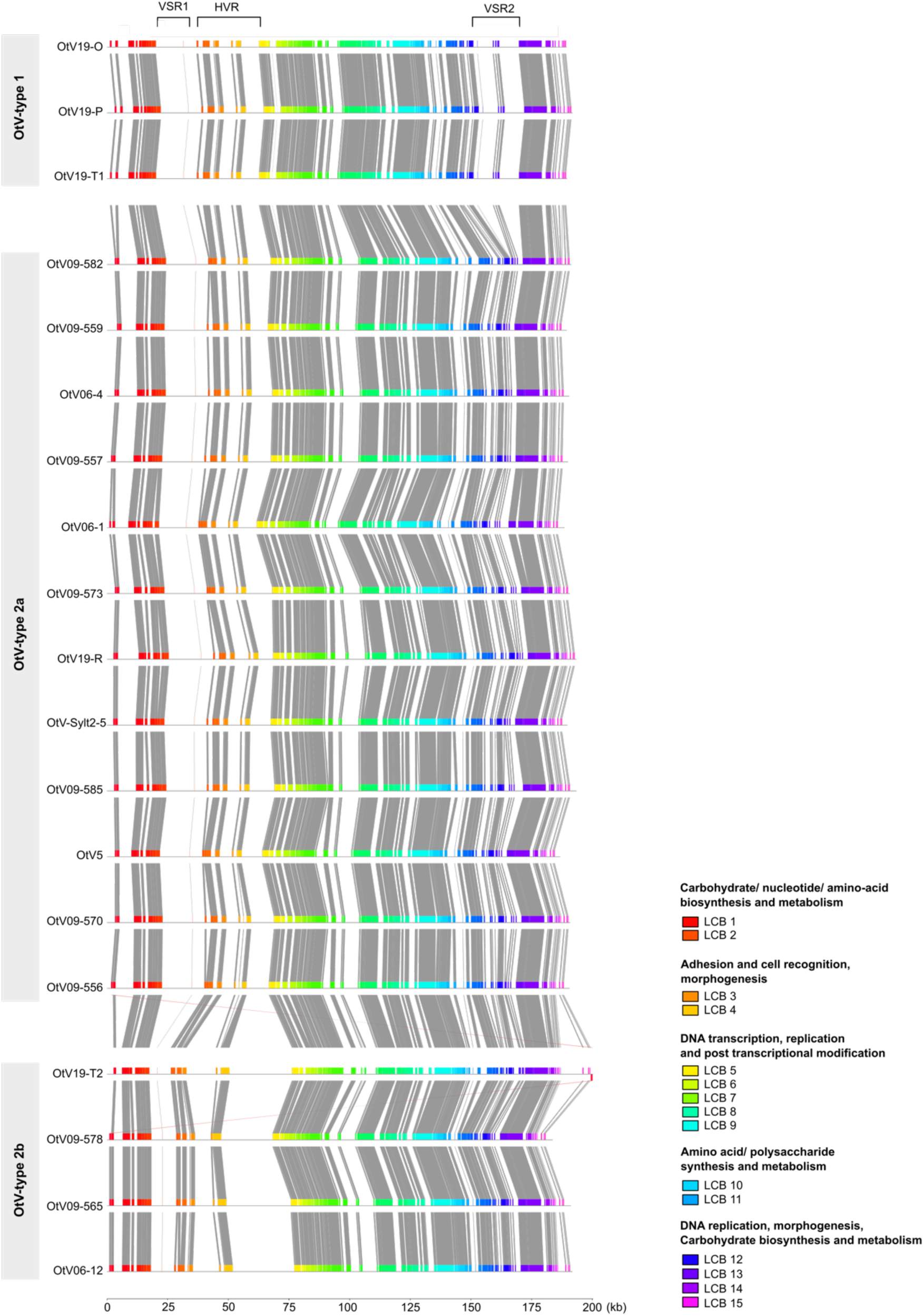
Comparison of the full-length genomes of *Ostreococcus tauri* viruses. Colored blocks correspond to conserved regions that are locally collinear and homologous between viral genomes. Such segments are referred to as Locally Collinear Blocks (LCBs) and connected by the lines and indicate commonly conserved regions between all genomes. The genomes are sorted by OtV-types (OtV-type 1, OtV-type 2a and OtV-type 2b). The scale is indicated in kilobase (kb). VSR: virus-specific region, HVR: hypervariable region.

Minor LCBs, not shared by all viruses, were observed at the genome extremities (Figure 2). These regions, called virus-specific regions (VSRs), harbor genes only present in certain OtV-types. The first VSR (VSR1), spanning about 20-kb, was present in all viruses except OtV-type 2b (Supplementary Figure 5a). Most of these genes had functions associated with carbohydrate metabolism, such as glycosyltransferase, galactotransferase, thiamine pyrophosphate enzyme, NAD-dependent epimerase/dehydratase, GDP-mannose 4,6 dehydratase and other genes more associated with fatty acids, amino-acids and secondary metabolism. The second VSR (VSR2) consisted of two separated blocks encoding a total of twenty-three genes, which were only found in the three virus genomes associated with the OtV-type 1 (OtV19-O, OtV19-P, OtV19-T1) (Supplementary Figure 5b). Among these genes, only seven (30%) encoded a known function, namely, DNA polymerase X family, staphylococcal nuclease homologue, mitochondrial 18-kDa protein, mannitol dehydrogenase, 2OG- Fe(II) oxygenase, AN1-like Zinc finger and deoxycytidylate deaminase. These genes are associated with nucleotide transport and metabolism, replication, recombination and repair, carbohydrate metabolism and other diverse functions. Interestingly, highly similar homologs of these genes were also identified in *Micromonas* viruses. Phylogenetic analysis of the mitochondrial 18-kDa protein indicated a recent lateral transfer of this gene between green algae and viruses (Supplementary Figure 6).

Finally, a large non-homologous region was interrupted LCB3 to LCB5 (from position ∼25 kb to ∼80 kb) (Figure 2). This region, hereby described as hypervariable (HVR), was characterized by complex rearrangements (i.e., deletions, duplications, acquisitions) of genes primarily associated with adhesion and cell recognition functions (Supplementary Figure 5a), as well as morphogenesis such as capsid proteins. A notable feature of this HVR was its expansion in OtV-type 2b viruses, which formed a faster diverging branch in the phylogenetic analysis (Supplementary Figure 4).

### The hypervariable region is a hotspot of host interaction genes

The HVR was characterized by a strong decrease in nucleotide sequence identity between viruses of different types and higher identity between the viruses of the same OtV-type, except for viruses of OtV-type 2b that showed large structural variations (Supplementary Figure 5a). The HVR was flanked by homologous core genes (from LCB3 and LCB5). Two capsid proteins delimited it on either side, and a third capsid protein was found in the central part, except for OtV06-12 and OtV19-T2 that encoded a fourth homolog (Figure 3).

**Figure 3.**
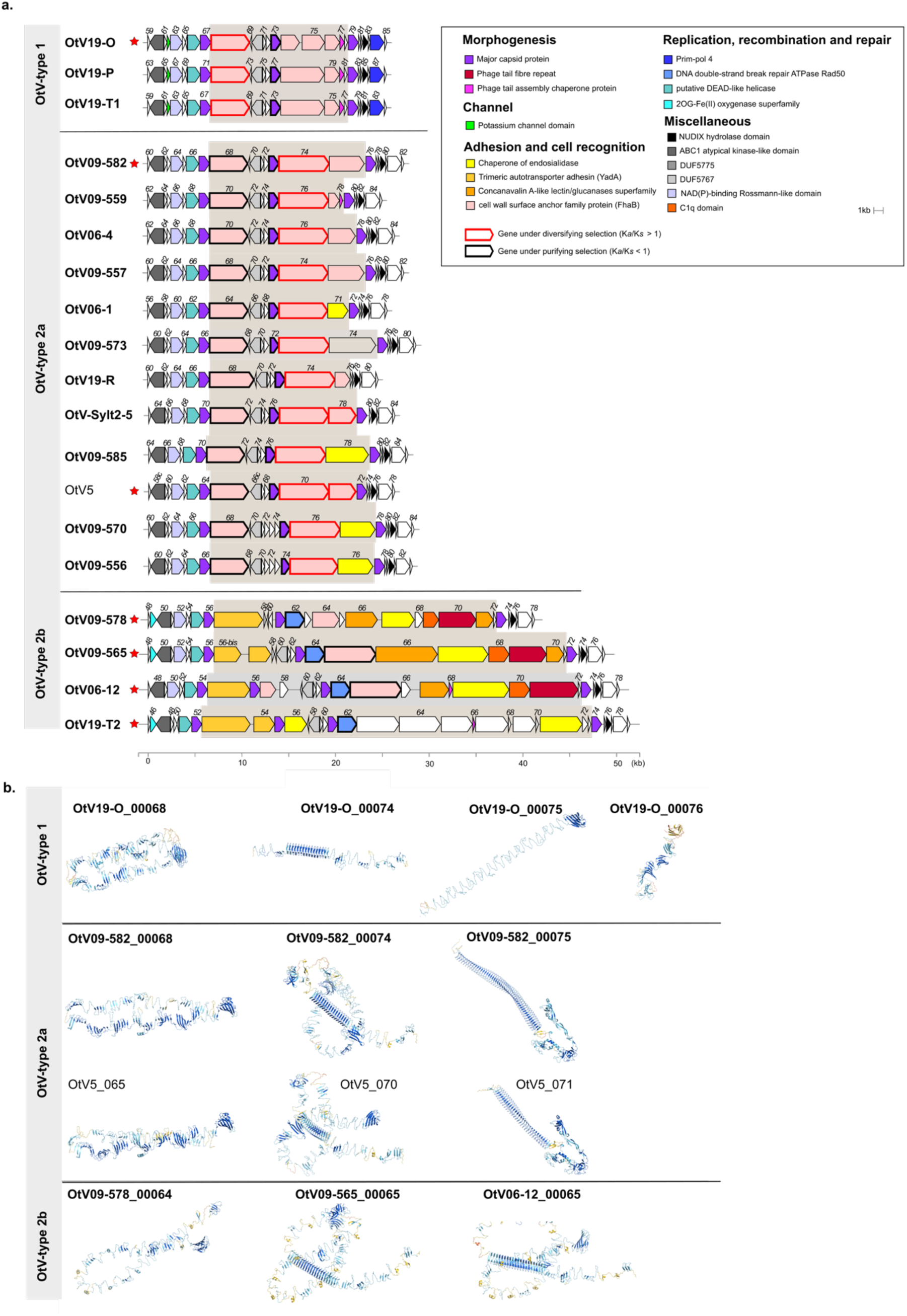
Description of the hypervariable region in OtV genomes. a) Genomic map comparing a partial sequence in which the hypervariable region was detected in the OtVs genomes. The hypervariable region is delimited by the box highlighted in gray. The OtV5 genome (unbolded) was used as a reference genome. Genomes were sorted according to their OtV-type based on their phylogenetic relationship and gene family composition (OtV-type 1, OtV-type 2a and OtV-type 2b). Red stars indicate viruses for which the search for gene function and/or conserved domain and structural domains was performed and then have been used as a reference for each OtV-type (Supplementary Table 6). Homologous genes were colored identically according to their functional categories. White open reading frames have no predicted function. Each open reading frame was numbered according to its associated locus tag. BLASTn comparisons between the OtV06-12, OtV09-578, OtV09-565, OtV19-T2, OtV5, OtV09-582, and OtV19-O genomic regions are shown in Supplementary Figure 4. The scale bar indicates the length of the genomic region in kilobase-pairs (kb). Genes under positive selection (*K_a_*/*K_s_* >1) are bolded and framed in red, while those under purifying selection (*K_a_*/*K_s_* <1) are bolded and framed in black. b) The three-dimensional (3D) of the cell wall surface anchor family protein (FhaB) associated with hypervariable regions in the OtVs genomes. The OtV06-12, OtV09-578, OtV09-565, OtV5, OtV09- 582, and OtV19-O were used as a reference for each type. Viruses isolated and sequenced in this study are bolded in black. A close-up view on β-folded segments of the FhaB protein is shown in Supplementary Figure 11. The full structural protein is shown in Supplementary Figure 11.

Upstream of the HVR were genes with functions associated with various biological functions, such as a DEAD-like helicase protein, a ABC1 atypical kinase-like domain-containing protein, a NAD(P)-binding Rossman-like domain-containing protein, and a 2OG-Fe(II) oxygenase. In addition, a putative potassium ion (K+) channel protein was also identified in the region upstream of the HVR. The K+ channel gene has been identified exclusively in OtV-type 1 (i.e., OtV19-P, OtV19-T1, OtV19-O, and OtV6) and in all *Micromonas* viruses including the newly characterized McV20-T (Supplementary 7). Furthermore, homologs were identified in the *Ectocarpus* virus and its host with a high identity (94.4%). Phylogenetic reconstruction of the K+ channel protein places prasinoviruses, chloroviruses, and TetV1 within a single monophyletic group with the filamentous brown alga *Ectocarpus siliculosus* and its virus (Supplementary Figure 7). While the evolutionary history of K+ channel protein seems difficult to assess between prasinoviruses and their hosts, our analysis indicated a recent lateral gene acquisition between *Ectocarpus siliculosus* virus and its host (Supplementary Figure 7). Interestingly, in OtV-type 2, a transmembrane protein with a distinct function was detected at the same genomic locus where the K+ channel gene was found in OtV-type 1 (Figure 3a) suggesting a likely evolutionary divergence of this gene between OtV-types.

The region downstream of the HVR consists of genes mainly associated with replication, recombination and DNA repair (i.e., prim-pol4 protein), as well as other functions such as NUDIX hydrolase domain protein, DUF5775 domain protein. Within the HVR, although most genes had unknown functions, most of the others were involved in cell recognition, adhesion and in morphogenesis. For OtV-type 1 and OtV-type 2a, these were predicted to be cell wall surface anchor family proteins (FhaB) and found in multiple copies. A phage tail assembly chaperone protein was identified in the OtV-type 1, as well as in one virus from OtV-type 2a (OtV09-559) and two viruses from OtV-type 2b (OtV06-12 and OtV19-T2). In addition, a chaperone endosialidase was predicted only in the OtV06-1, OtV09-585, OtV09-570 and OtV09-556 genomes. A remarkable feature of the HVR is its substantial expansion in the OtV-type 2b, which is more than three times longer than that of other viruses (Figure 3). For example, the average length is 15-kb in OtV-type 1 and OtV-type 2a, compared to about 40-kb in OtV-type 2b (Figure 3). This difference in size was notably associated with the acquisition of unique genes that are most likely involved in host interaction and recognition. These genes encode phage tail fiber proteins, concanavalin A-like lectin/glucanase superfamily proteins, and trimeric autotransporter adhesins (YadA). In addition, a gene with a putative C-terminal Rad50 domain protein, located near clusters of variable host specificity and interaction genes, was only found in the OtV-type 2b virus genomes (Figure 3). In eukaryotes, the Rad50 protein is associated with the Mre11 protein (the bacterial homolog of the SbcD/SbcC system (Pandey et al. 2018)) which are involved in repairing DNA double-strand breaks (Lammens et al. 2011; Storvik & Foster 2011). The structure of this protein in OtV-type 2b viruses was predicted with low confidence (pTM = 0.29) to fold as a long alpha-helix coiled-coil domain and two mixed beta-strands in N- and C-terminal. Foldseek search revealed no obvious function and did not detect any similarity with Mre11. Tentative pairwise alignment with *eg.* 5F3W and 4YKE Mre11 experimental structures failed as well. However, due to the low homology of Rad50 protein with other organisms and the lack of homology for a Mre11-like domain, it remains challenging to draw any conclusions about the function of the gene and its association with the large structural variations specific to OtV-type 2b.

To identify molecular signatures associated with evolutionary processes in host–virus interactions, the ratios (ω) of non-synonymous (Ka) and synonymous (Ks) substitution rates were estimated in each OtV-type across different alignments for genes within the HVR with nucleotide of sequence >80% of identity (Figure 3; Supplementary Table 5). A total of seven distinct groups of homologous genes (orthogroups) could be tested. Of these seven orthogroups, five showed a low ratio across five different models tested. This concerns a capsid protein central to HVR for both OtV-type 1 and OtV-type 2a groups (an average of ω=0.02 and p-value=2E-105), a gene coding for a rad50 protein in the OtV-type 2b group (an average of ω=0.21 and a p-value=3E-68) and one of the cell wall anchor surface familly proteins (FhaB) in the OtV-type 2a group (an average of ω=0.1 and a p-value=3E-16). Ka/Ks ratios below one are indicative of purifying (negative) selection, implying that these proteins are functionally constrained and likely play critical roles in the viral infection cycle. Interestingly, two orthogroups, both coding for putative FhaB proteins, showed high Ka/Ks ratios in the OtV-type 1 and the OtV-type 2a groups. While for OtV-type 1, Ka/Ks ratios vary between models, the NG model suggests significant positive selection, with an average of ω=1.42 and a p-value=0.008. For OtV-type 2a, Ka/Ks ratios were consistently >1 across all models, with highly significant p-values (average of ω=1.3 and a p-value=3E-155), indicating strong diversifying (positive) selection. These results suggest these proteins may be subject to rapid adaptive evolution, possibly in response to host interactions and may therefore contribute to the expansion of the host range.

To further investigate the putative functions of genes associated to the HVR, 124 protein sequences, from all genomes in the OtV-type 2b, the OtV19-O (OtV-type 1), and two genomes from OtV-type 2a (OtV09-582 and OtV5), were submitted to AlphaFold3 for the computational prediction of their three-dimensional structures (Abramson et al. 2024). The length of the sequences ranged between fifty-eight and 2,188 amino acids. All targeted proteins yielded predictions with overall accuracy reported by template modeling scores (pTM) ranging from 0.23 to 0.93. Confident high-quality predictions were made for thirty-seven protein models (pTM≥0.80), twenty-one had intermediate confidence (0.60≤pTM≥0.80) while sixty-six models were not sufficiently reliable to obtain structural predictions (0.60≤pTM). We chose fifty-eight of the 124 models for submission to a protein structural similarity search with Foldseek (van Kempen et al. 2024). Principal hits were searched in the structural databases AlphaFold/Proteome, AlphaFold/Swiss-Prot, AlphaFold/UniProt50, and PDB100, confirming the presence of peptidase S74, ABC1 domain-containing protein, autotransporters, glycosyl- or carbohydrate-related proteins, and phage-tail proteins. Detailed analysis on thirteen FhaB common to the HVR revealed elongated β folds (Figure 3b). The viral FhaB homologs (OtV06-12- 00065, OtV09-565-00065, OtV19-O-00074, OtV09-582-00074, and OtV5-070) were predicted to form a right-handed β-helix structure consisting of 42-63 strands, with three faces each made up of parallel β-strands. Overall, they form a β-3-solenoid (CATH ID: 2.160.20). The two homologs in OtV09-582 and the OtV5 (i.e., OtV09-582-00075 and OtV5-071) were predicted to form an elongated 30-48 strand single parallel β-sheet folding as a right-handed helical solenoid and respectively spanned 21.2 nm and 13.6 nm in their longest dimension. Both fibrillar architectures are therefore likely to generate extra-virion projections and facilitate intermolecular recognition.

### OtV-type 2b viruses display the broadest host range

A cross-infectivity assay was carried out on all viruses studied in this work against a panel of nine *O. tauri* strains by spotting viruses onto hosts grown on soft agarose plates. Note that both OtV19- T1 and OtV19-T2 viruses originated from the same lysate (OtV19-T) and therefore did not allow us to study the host range of these viruses separately.

Three different viral infection patterns were observed: i) No lysis, which was characterized by the absence of plaques, ii) strong lysis, was defined by large and well-rounded plaques and iii) lower lysis, which gave smaller and more turbid plaques (Figure 4A). As expected, only *Ostreococcus* strains were susceptible to infection by OtVs (Figure 4B). The two *Micromonas* and *Bathycoccus* strains showed no lysis, indicating a specificity of the genus *Ostreococcus* of these viruses. All lysates infected at least one of the *Ostreococcus tauri* strains tested, while OtV-Sylt2-5 and OtV09-573 were also able to infect *O. mediterraneus* strain RCC789. Our analysis revealed that the *O. tauri*-OtV cross-infection assay exhibited a "nested" or hierarchical structure. In this structure, the viral host range was on a continuum from specialized viruses infecting a narrow range of host strains to generalist viruses infecting a broader range of hosts. The OtV5, OtV06-1 and OtV19-O viruses displayed the narrowest host range (i.e., specialist), infecting only the most susceptible strain, *O. tauri* RCC4221. In contrast, the OtV09-578, OtV06-12, OtV09-565 viruses, which were associated with the OtV-type 2b group and whose hypervariable region was characterized, were viruses with the broadest host range (i.e., generalists), infecting all the strains tested (Figure 4B).

**Figure 4.**
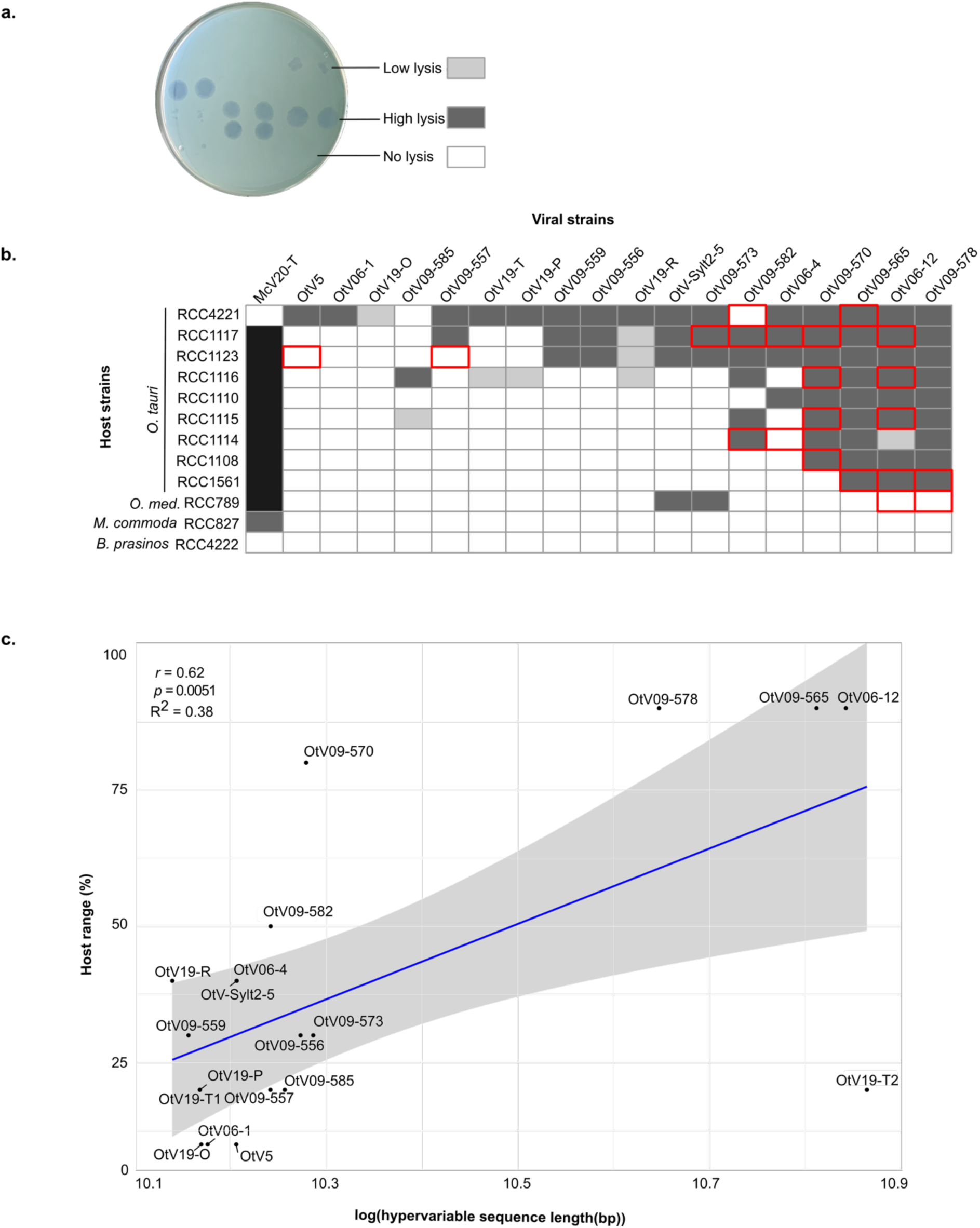
Host–virus infection patterns. a) Example of plaque assays for different viral infections showing low, high and no lysis for a given phytoplankton host strain. b) Cross-infectivity assays of OtVs against a panel of *Ostreococcus* sp. strains (*O. tauri*, *O. mediterraneus*) and two representative species of the most closely related genera within the Mamiellophyceae class (*Micromonas commoda* and *Bathycoccus prasinos*). Dark gray box corresponds to no lysis observed, white indicates lysis was observed three days post infection and light gray indicates lower lysis, where lysis plaques were smaller or more turbid. Black indicates not tested. Note that phenotypic changes observed compared to the data of Clerissi *et al*. 2012 were framed around the boxes in red. The host range of McV-20T was only tested on the strains *Micromonas commoda* RCC827, *Ostreoccocus tauri* RCC4221 and *Bathycoccus prasinos* RCC4222. c) Linear correlation between the host range and the hypervariable sequence length of OtVs. The host range is represented in percentage i.e., The number of successful lysis divided by the total number of *Ostreococcus* strains tested in the experiment (i.e., 10 strains). The x-axis is log-transformed. Confidence interval is indicated in light gray. *r*: Pearson correlation coefficient. *p*: *p*-value and R^2^: coefficient of determination.

Interestingly, 67% of viruses (8 out 12 viruses) showed changes in infection pattern in comparison to previous analysis (Clerissi et al. 2012). The switch from non-infectivity to infectivity was the most reported phenotypic change (15 out of 20 — 75%), rather than the shift from infectivity to non-infectivity (5 out of 20 — 25%) (Figure 4B). In addition, those changes were mostly observed for the two most generalist viruses (i.e., OtV06-12 and OtV09-570) and for one of the host strains most susceptible to infection (*O. tauri* RCC1117 strain). Our analysis revealed a statistically significant, although moderate, positive correlation between the host range and the HVR length (*p* = 0.0051, *r* = 0.62, R² = 0.38) (Figure 4C), implicating these gene expansions as the genetic determinants of increased host range. Nonetheless, two OtV strains stood out as clear outliers in terms of this association, namely, OtV09-570, which had no obvious expansion in HVR size but an apparently broad host range; and OtV19-T2, which had a large HVR expansion but narrow host range. As the variability in host-range over time particularly affected OtV09-570, it suggested this strain may be given to rapid variation in host range with time or be phenotypically variable. This variability in host range clearly complicated a direct association between genotype and host range.

### The O. tauri-prasinovirus dynamics are highly variable between strains

To compare the dynamics of infection between viral strains of the same phylogenetic clade with the aim to better link viral genetic features to infection traits, four *O. tauri* strains (RCC4221, RCC1123, RCC1116 and RCC1108) with a range of patterns of susceptibility to OtVs (Figure 4) were challenged with the sixteen OtVs of type 2 (Figure 5). Overall, the infection dynamics differed between the different *O. tauri* strain populations showing strain-specific responses to infection by OtV strains. As expected, the *O. tauri* RCC4221 was the most susceptible strain to all the viruses tested showing lysis in response to the same virus strains as in the cross-infection spot-based assay (Figure 4). In addition, the lysis dynamics of *O. tauri* RCC4221 in response to infection by OtV strains occurred along a continuum in terms of time to lysis, speed of lysis and mortality. By contrast, *O. tauri* RCC1108, which was the least susceptible strain (Figure 4), showed strong lysis in response to two OtVs (OtV09-565 and OtV09-578) in liquid culture (Figure 5a and Supplementary Figure 8). However, RCC1108 was not lysed by two additional viruses, OtV09-570 and OtV06-12, as in the spot assay (Figure 4). Similarly, RCC1116 was lysed in liquid culture by four viruses (i.e., OtV09-565, OtV09-578, OtV09-582, OtV09-585) but was not visibly lysed by an additional four viruses (i.e., OtV06-12, OtV09-570, OtV19-T, OtV19-R) that produced plaques in Figure 4. Intriguingly, RCC1123 showed the inverse case with lysis occurring from infection by OtV09-557 in liquid medium (Figure 5A and Supplementary Figure 8), but with no plaques in the spot assay (Figure 4). Host-strain specific responses were evident in the comparison of the lysis dynamics of the most generalist virus strains, OtV09-565 and OtV09-578, which showed the degree of mortality varied with each host strain. For example, RCC1116 lysed relatively slowly in response to infection by these two viruses, whereas it showed a higher degree of mortality to strains OtV09-582, OtV09-585. Altogether, this revealed that successful lytic viral infection can indeed depend on the growth conditions, in this case if the growth medium was semi-solid or liquid indicating a factor strongly influencing the variability in host range. The variation in host range between the two assays was most pronounced for the strain OtV09-570, which lysed only RCC4221 instead of all four tested strains as expected from the plaque assay, again indicating it as the most variable in host range.

**Figure 5.**
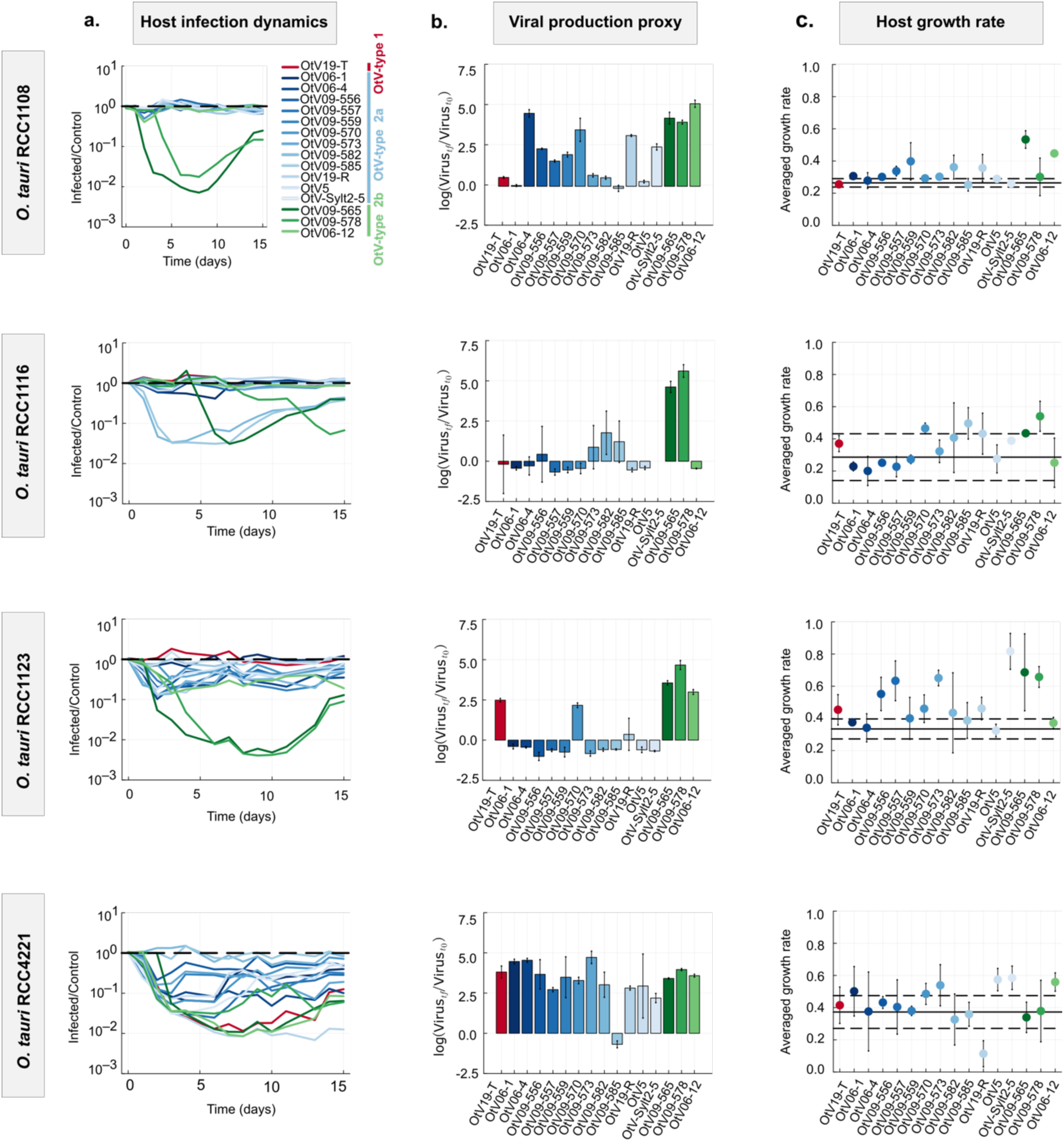
Infection dynamics of *Ostreococcus tauri* strains and the OtVs. The four *O. tauri* strains tested (RCC1108, RCC1116, RCC1108 and RCC4221) were sorted in rows and for each host the following parameters were measured: a) Averaged host dynamics (*n* = 3) plotted as the cell fluorescence ratio in infected *vs.* non-infected (control) cultures. The black dashed line corresponds to the baseline cell density of the control. Colored lines below the dashed line indicate infected cultures with a decrease in cell density relative to the uninfected controls. b) Proxy of virus production calculated as the log-transformed ratio of virus concentration at the final time *vs.* inoculation time. c) Averaged population growth rate (in days) of the cells in the recovery phase. Error bars represent the standard deviation to the mean (*n* = 3). Solid and dashed black lines represent the averaged growth rate and standard deviations, respectively, in the control culture. The growth rate has been estimated as the slope of the linear regression of the log-transformed cell concentration during the exponential re-growth in the recovery phase.

We measured viral particle concentration at the end of the 15-day time course and observed that the host lysis dynamics were not well-associated with our proxy of viral production (Figure 5b). Clear virus production could only be associated with lysis of the most susceptible strain RCC4221, and with strong lysis by the most generalist strains OtV09-565 and OtV09-578, which allowed us to infer these host–virus interactions resulted from productive infections. However, there were multiple apparent cases of virus production that were not associated with cell lysis. The clearest example was the *O. tauri* RCC1108 strain, which produced OtV06-4 viral particles at a rate of five, similar to the OtV09-565 virus, but with no host mortality. Another notable example is the *O. tauri* RCC1123 strain, in which the OtV19-T virus produced 2.5 viral particles, while no host lysis was detected.

### Prasinoviruses show distinct morphologies

Morphological investigations using transmission electron microscopy (TEM) were carried out on the new viruses isolated in this study (i.e., OtV19-P, OtV19-O, OtV19-T, OtV19-R and OtV-Sylt2- 5), as well as on the viruses associated with OtV-type 2b (i.e., OtV06-12, OtV09-578 and OtV09-565), in comparison with the OtV5 virus previously characterized (Derelle et al. 2008).

Overall, the OtVs shared a common morphology; they produced non-enveloped virions with an icosahedral capsid having an averaged diameter of 140 ± 8 nm (Figure 6a). Furthermore, notable phenotypes were observed for the two viruses OtV09-565 and OtV19-T. The OtV19-T lysate, in which two distinct genotypes were characterized in this study, consisted of two morphotypes (Figure 6a; Supplementary Figure 9a). The first morphotype was comparable to that observed for the majority of OtVs, i.e., non-enveloped with an icosahedral capsid, with a diameter of 139 ± 10 nm. By contrast, the second morphotype appeared "squashed", with the flattening of a vertex and the widening along the diameter (148 ± 8 nm). Such changes might be the result of genetic and/or structural constraints that affected capsid protein arrangement, especially under specific environmental conditions. It is tempting to speculate that the two morphologies of OtV19-T corresponded to the two different genotypes. Finally, a striking new morphological feature was observed in OtV09-565. This virus produced virions with an icosahedral capsid (averaged diameter of 140 ± 9 nm) to which was connected an elongated and flexible structure of 154 ± 46 nm and 21 ± 5 nm in length and width, respectively (Figure 6a and b). This structure appeared to be tail-like and superficially similar to those of viruses in the *Caudovirales*. It emerged from one vertex, but no other structures, such as spikes or a base plate, were observed. On the other hand, the lysate consisted of a mixture of tailed and non-tailed virions, with a slightly higher proportion of the tailed particles (Figure 6c). Free tails were also observed next to non-tailed particles, indicating the lability of this structure (Supplementary Figure 9b). Moreover, no significant difference in capsid size was detected between the two types of virions, supporting the hypothesis of a single viral population (Figure 6b).

**Figure 6.**
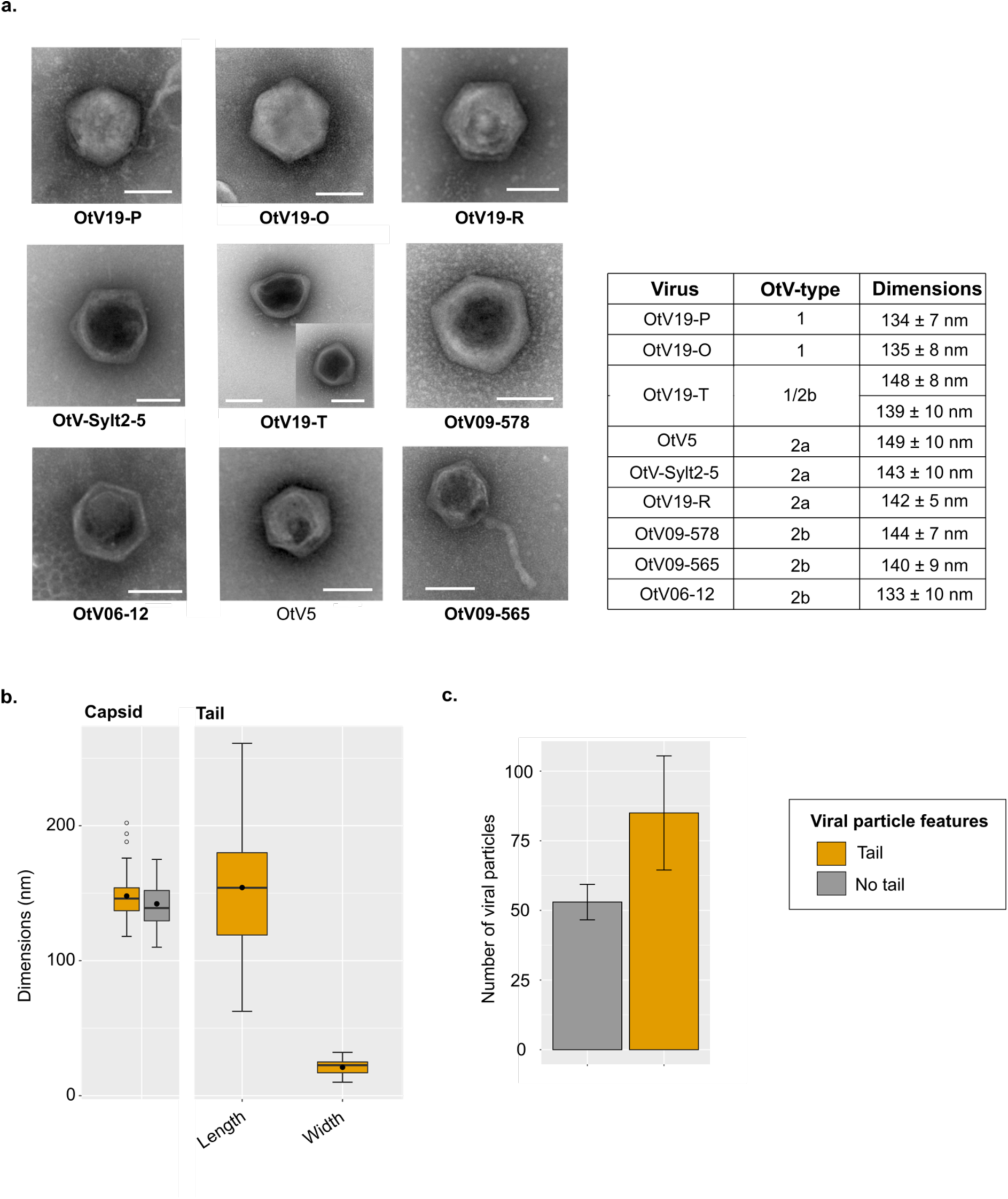
Morphological characterization of *Ostreococcus tauri* viruses. a) Transmission electron micrograph of OtV particles. The viruses isolated from the Pacific Ocean (OtV19-O, OtV19-P, OtV19-T, OtV19-R) as well as the North Sea virus (OtV-Sylt2-5) were compared to the Mediterranean viruses associated to OtV-type 2b than OtV5 (OtV-type 1). Viruses newly characterized in this study are bolded. The scale bar corresponds to 100 nm. The dimensions are shown as the average and standard deviations of the capsid dimensions (width). b) The capsid size of the tailed-virion was compared to the non-tailed one, and the length and width of the tailed-viral particles were reported. c) Bar plot showing the number of tailed-virion (orange) or non-tailed (gray) in a lysate obtained three days post-infection.

Finally, to further investigate the biological function and potential role of the tail-like structure of OtV09-565 during infection, thin sections of both uninfected and infected algal cultures were examined using TEM. For uninfected conditions, images revealed healthy cells, as expected, with intact organelles such as chloroplast, nucleus, mitochondrion, cytoplasm, and starch grain (Supplementary Figure 10). In infected cells, a viral particle was observed near a cell 5 minutes after infection. While a thin structure connecting the capsid and the cell surface was observed during the attachment phase (at 30 min), no tail-like appendage associated to the capsid could be identified in this case. Intracellular virions were first observed in the cytoplasm at 4 hours post-infection (hpi). By 12 hpi, the cell became packed with viral particles, ready for lysis to release the newly formed virions. Finally, at 72 hpi, the cells are completely lysed with viral particles in suspension in the medium (Supplementary Figure 10). The tail-like structure was not observed in any stage during assembly or release.

## Discussion

In this study, we sought to expand our knowledge about the diversity of the *Prasinovirus* genus by isolating new viruses from distant locations around the world. The sequencing of eighteen new complete OtV genomes has shed light on the diversification and evolution of these viruses and the genetic signatures associated with their host range. To date, prasinovirus genomes have been observed with a high degree of conservation of the order of homologous genes (Moreau et al. 2010). In this work, we describe an organization of OtV genomes modularly defined with both synthetic and non-syntenic regions. A similar pattern was observed in *Ostreococcus lucimarinus* virus (OlV) genomes, where a central and large DNA fragment was inverted, distinguishing two subgroups of OlV (Derelle et al. 2015). Here, we defined three distinct new OtV clades, classified as OtV-type 1, OtV-type 2a and OtV- type 2b which showed distinct evolutionary relationships. Notably, OtV-type 1, relatively divergent from other OtVs, forms a subclade within the *Micromonas* viruses clade. Furthermore, approximately 60% of the OtV-type 1 clade-specific proteins were homologs in the genomes of *Micromonas* virus, which were absent from all other *Ostreococcus* viruses. Our results were consistent with the previous work of Thomy et al. using a targeted enrichment approach in the South China Sea (Thomy et al. 2024). The authors identified a new viral clade, termed clade III, consisting of a set of *Micromonas* and *Ostreococcus* viruses, including the previously sequenced OtV6 genome. This clade branches into the *Micromonas* virus clade. Here, our work further supports the idea of host switching between genera and the hypothesis of host range expansion, although these viruses failed to infect the *Micromonas commoda* RCC827 strain. We also suggest the possibility that these viruses evolved from a common ancestor infecting the hosts *Micromonas* and *Ostreococcus* or an intermediate lineage. These results raise several questions about the evolution and ecology of these viruses. Could these viruses expand their host range by infecting the same host through recombination events and horizontal gene transfer? Although no evidence of homologous recombination events was found in the OtV genomes in this study, we were able to describe the horizontal transfer of a gene encoding the 18 kDa mitochondrial protein from *Ostreococcus* and *Micromonas* strains to OtV-type 1 viruses and some *Micromonas* viruses. This result suggests that the acquisition of this gene occurred independently in both lineages or that a common ancestor acquired it from a host before diverging into two separate lineages. Although the exact role of the 18 kDa mitochondrial protein in algae is not yet fully understood, in animals this receptor is recognized as crucial for cellular respiration, steroid hormone production, cell death, and immune response (Papadopoulos et al. 2006). Hence, it is likely that the 18 kDa protein plays a key role in host metabolism, potentially benefiting viral infection. This strongly supports the hypothesis that OtV-type 1 have adapted both to their hosts and to the environmental conditions. In addition, most of the clade-specific genes, including the 18 kDa protein, were found in a cluster, appearing in genomic islands or, as described in this study, virus-specific regions (VSRs).

The adaptability of viruses to extend their host range is driven by genetic changes such as mutations, recombinations, and rearrangements mainly targeting genes involved in attachment to a receptor at the host cell surface (Ferris et al. 2007; Zhao et al. 2019). In bacteriophages, such mutations have been found in host recognition modules encoding tail fibers, chaperones and baseplates (Hashemolhosseini et al. 1994; Tétart et al. 1996; Kizziah et al. 2020; Gonzalez-Serrano et al. 2023). In contrast, the genetic basis of host-range evolution in eukaryotic dsDNA viruses is poorly understood. In this study, we highlighted a HVR in the OtV genomes that is characterized by numerous mutations in genes encoding surface proteins and/or receptor-binding domains involved in viral recognition and attachment to the host cell. Furthermore, this high genetic plasticity was mainly detected in OtV-type 2b, the group of viruses with the broadest host range. We propose that this region acts as a hotspot for genomic recombination and diversification. This process is driven by strong coevolutionary arms-race dynamics, in which viruses continuously adapt to overcome host defenses and hosts evolve to evade viral infection. Such dynamics parallel those extensively documented in bacteriophage-bacteria systems, where host recognition genes undergo rapid diversification (Comeau et al. 2007; Labrie et al. 2010).

Additionally, our data revealed that viruses from geographically distinct locations can infect identical host strains. One possible explanation involves an alternative infection pathway that enables these viruses to target the same host strain without necessarily broadening their host range. Notably, a specific genomic region identified in this study as VSR1 was exclusively shared between OtV-type 1 and OtV-type 2a, two virus groups originating from different geographical regions. This region encodes genes involved in fatty-acid, amino-acid and secondary metabolisms as well as carbohydrate metabolism including glycosyltransferases. In contrast, only glycosyltransferases with low sequence homology with other groups were found in OtV-type 2b viruses from this same region. The carbohydrate biosynthetic enzymes, including glycosyltransferases, have been documented in giant viruses (Speciale et al. 2022). Glycosylation-related genes encoded by viruses allow them to become partially or completely host-independent by modifying their own glycoproteins. This feature was thought to be a viral strategy for evading cellular immunity by mimicking host glycans. In addition, the glycosylation of capsid proteins may also influence host cell attachment and entry, playing a potential role in host specificity. To date, glycosylation of the chlorovirus capsid protein has been the most widely studied of the giant viruses (I. N. Wang et al. 1993; Graves et al. 2001; Nandhagopal et al. 2002; Van Etten et al. 2010; Xiang et al. 2010). However, although the exact mechanism of glycosylation in prasinoviruses is still poorly understood, we propose that this region might be an alternative pathway encoding the glycosylation of the viral capsid in determining host specificity.

Furthermore, our analysis revealed variability in the host range, which particularly affected OtV09-570. This suggests that this virus may undergo rapid changes in host range over time or exhibit phenotypic variability. This variability in host range clearly complicated a direct association between genotype and host range. Given this pattern of HVR variability and variability in host range, we hypothesized that the genetic determinants of the host range likely involved a complex mechanism of multiple genetic features interacting specifically with the host strain genetics. Moreover, given the variation over time in our spot assay, host range may be evolving rapidly over time, or it may be a variable trait when measured with this assay. Also, host range may be sensitive to variations in other, yet unidentified factors, such as variations in abiotic factors. Beyond the host range, analyses of infection dynamics have revealed substantial temporal variability. Initially, the lack of correspondence between lysis and viral production was due to a limitation of the experimental setup, in which we could only measure virus concentration at the end of the experiment. This meant that we could not observe virus dynamics, such as virus decay, during the 15-day time course. This phenomenon could also be explained by i) heterogeneity in the host populations, such as virus production from a subpopulation of susceptible cells in a majority virus-resistant population; or ii) a chronic infection in which viral particles are produced by a budding mechanism without cell lysis. The former case was observed in resistant populations of *O. tauri* and *O. mediterraneus* (Yau et al. 2016, 2020), while the latter hypothesis was previously proposed by Thomas et al. for the *O. tauri* strain and the OtV5 virus as a mechanism of viral resistance (Thomas et al. 2011).

The acquisition of the resistance, or cell recovery, is defined as the phase in which the concentration of viable cells re-increases after a lysis phase. This phenomenon has been previously described in Mamiellophyceae (Thomas et al. 2011; Yau et al. 2018). In this work, we observed that resistance occurred in all cross-infections (30 crosses) except for two (i.e., OtV19-R on RCC4221, OtV09-578 on RCC1116) in the 15-day time frame of the experiment (Figure 5c and Supplementary Figure 8). This indicated that development of resistance was a highly consistent trait irrespective of the host–virus strain interactions. In addition, all recovered cultures showed a population growth rate equal to or higher than that of the uninfected culture (control) (Figure 5c), suggesting that the development of resistance had no detectable “cost of resistance” trade-off (B. j. m. Bohannan & Lenski 2000; B. J. M. Bohannan et al. 2002) visible in the population growth rate. Overall, these findings showed that the *O. tauri*-prasinovirus system has a high variability in the infection dynamics, which was specific to the host and virus strain combination, and the host recovery response was a highly consistent trait in the *O. tauri* strains tested.

By integrating computational and experimental approaches, our study offers comprehensive insights into genetic diversity of *Ostreococcus tauri* viruses and, for the first time, describes a form of viral plasticity potentially driven by antagonistic coevolution with host organisms. In addition, we report novel morphological features that may play a key role in host–virus interactions. Overall, these data provide a better understanding of how marine viruses that infect eukaryotes adapt and evolve in response to changing environmental conditions and pressures from the host immune system in the ocean.

## Material and Methods

### Culture of host algae and viruses

All algal cultures and viruses (Supplementary Figure 1) were grown in liquid L1 medium (NCMA, Bigelow Laboratory for Ocean Sciences, USA) made with autoclaved offshore seawater (MOLA station: 42°27’11”N, 3°8’42” E), diluted with Milli-Q water to give a final salinity of 35 PSU, and filter sterilized through 0.22 µm polyethersulfone (PES) filters. The cultures were maintained at 15°C using a photoperiod cycle of 12 h: 12 h light: dark (50 µmol photon m^-2^ s^-1^ white light).

### Isolation of new viruses and host-virus specificity tests

Viruses from the Mediterranean Sea had been previously isolated between 2006 and 2009 (Clerissi et al. 2012). One virus from the North Sea and six viruses from the Pacific Ocean were isolated in this study. These viruses were isolated from half a liter of seawater collected from each marine station site (Supplementary Table 1) that was filtered through a 0.45-µm PES membrane filter (SARSTEDT reference; 83.1826), concentrated ten times by tangential flow filtration (100 kDa PES Vivaflow200, Sartorius, Germany) and then clonal viruses were procured by plating for individual plaques in exponentially growing *O. tauri* RCC4221 cultures, as described previously (Derelle et al. 2008). We considered that the virus strains were clonal after three successive infections of the host strain by the viruses from one lysis plaque. Fresh viral lysate was filtered through 0.22-µm or 0.45-µm pore-size filters to remove cellular debris and stored at 4°C in the dark. For host-virus specificity tests, a panel of Mamiellophyceae strains from the Roscoff Culture Collection (www.roscoff-culture-collection.org) (*Ostreococcus tauri* RCC4221, RCC1123, RCC1117, RCC1116, RCC1110, RCC1115, RCC1108, RCC1114, RCC1561; *O. mediterraneus* RCC789; *M. commoda* RCC827; and *B. prasinos* RCC4222) were screened for their susceptibility to the viruses analyzed in this study. Algal strains in exponential growth were incorporated into semi-solid L1 medium by addition of 1.5% molten agarose held at 58°C in a water bath (0.15% final concentration) and poured into Petri dishes (9 cm diameter). Three microliters of each viral lysate were deposited in duplicate on the surface of the semi-solid medium for each algal strain, incubated at 20°C using a photoperiod of 12 h: 12 h light: dark cycle (50 µmol photon m^-2^ s^-1^ white light), and the susceptibility to viral lysis was evaluated by the appearance of plaques visualized after 7 days.

### Host-virus infection dynamics

Cross-infection assays were performed in a 96-well cell culture plate (200 µL final volume) in triplicate using a host cell concentration of 10⁶ cells/mL and 10^7^ viral-like particles/mL of virus (virus to host ratio of 10). Viral particle concentration was measured at the final time point and compared to known concentration at time zero. The plate was incubated at 20°C using a photoperiod cycle of 12 h: 12 h light: dark (50 µmol photon m^-2^ s^-1^ white light). The growth of the host cells was monitored using chlorophyll *a* autofluorescence (excitation/emission: 430/670 nm) in a Victor Nivo microplate reader over 15 days. To estimate the number of virus-like particles produced during infection, all the viral lysates were collected, and the concentration of viruses was measured by flow cytometry at the end of the 15 days of the experiment, as described by Brussaard (Brussaard 2004). Briefly, viruses were fixed with 0.5% glutaraldehyde (25% electron microscopy grade) for 15 min and flash frozen at stored at - 80°C until measured. Virus particles were thawed, diluted in TE buffer (pH=8.2, 10 mM Tris–HCl, 1 mM EDTA filtered through 0.02 µm pore-size filters (Anotop syringe filter, Whatman) marked with SYBR-Green (10^-5^ dilution of stock, Thermo Fisher Scientific) by heating to 80°C in the dark for 10 mins. Virus particles were enumerated in the CytoFLEX flow cytometer (Beckman Coulter) gated on green fluorescence *vs*. side scatter. *Ostreococcus* growth rate in the re-growth exponential phase was calculated as the slope of log(concentration) regression. Re-growth exponential phase corresponds to the phase where the *Ostreococcus* growth rate becomes positive after the culture reaches its minimum concentration and before reaching the saturation phase again. If necessary, we removed points at the border of the re-growth exponential phase to ensure estimating the maximum growth rate using the best possible regression fit that minimizes the error between the model and the data.

### Transmission electron microscopy

Observation of viral particles in negative staining was performed on copper grids coated with Formvar film and carbon film (FCF200-Cu - EMS), rendered hydrophilic by effluviation (20 seconds of air glow discharge,-ve, GloQube® Plus Glow Discharge System). The grid was inverted onto a drop of fresh 0.45 μm-filtered viral lysate for 30 minutes to allow viruses to adhere to the grid support film. After removal of excess sample by adsorption onto filter paper, the grid was laid on a drop of 2% aqueous uranyl acetate solution for 45 seconds. After absorbing the excess dye with filter paper, the grid was air dried. The observation was performed on a Hitachi 7500 TEM.

For thin-section observation by TEM, non-infected (control) and infected *Ostreococcus tauri* cells (infected with OtV09-565) were prepared following the protocols of Chretiennot-Dinet et al. (Chrétiennot-Dinet et al. 1995) and Derelle et al. (Derelle et al. 2008). A concentration of 15 × 10⁶ cells/mL, contained in 50 mL aliquots of samples pre-fixed with 1% glutaraldehyde (Electron Microscopy Sciences), was collected by centrifugation at 2500 × g for 30 minutes. The pelleted cells were then quickly mixed with 40 µL of low melting point agarose (Sigma) at 37°C using a micromanipulator (SMI, Emeryville, CA, USA). Once the agarose solidified, the agar block containing the cells was fixed for 2 hours at 4°C in 2.5% glutaraldehyde prepared with one volume of 0.4 M cacodylate buffer and two volumes of culture medium (L1 medium) and washed three times for 15 minutes each in one volume of 0.4 M cacodylate buffer and one volume of culture medium. Post-fixation was carried out in 1% osmium tetroxide (OsO₄) (Electron Microscopy Sciences) in 0.2 M cacodylate buffer for 1 hour. After two washes in the same buffer, the small agar blocks were gradually dehydrated in ethanol (70%, 95%, 100%), then infiltrated and embedded in Epon 812 resin (Electron Microscopy Sciences). Ultrathin sections (80 nm) were obtained using an ultramicrotome (Leica Ultracut R) and examined on a Hitachi 7500 TEM after staining with uranyl acetate and lead citrate. The resulting images were processed using ImageJ software.

### Partial DNA polymerase B gene amplification and sequencing

We determined that the newly isolated viruses were prasinoviruses based on PCR screening of the partial *polB* gene, which is a viral marker gene, using the primer sets in Clerissi *et al*. 2014 (Clerissi et al. 2014). PCR reactions were done as follows: 2 µL of fresh viral lysate was added to a 48 µL of reaction mixture containing 1X PCR buffer, 1 mM MgCl₂, 0.2 mM deoxyribonucleoside triphosphate (dNTP) mix, 0.5 µM of each primer, 0.1 mg.m^-1^ BSA and 1.25 U of Taq DNA polymerase (GOTaq® G2 Flexi M7805, Promega). PCR was performed in a Mastercycler Nexus system (Eppendorf) using an initial denaturation at 95°C (3 min), 38 rounds of denaturation at 95°C (30 s), annealing at 49°C (60 s), extension at 72°C (3 min) and final extension at 72°C (4 min). PCR products were loaded into a 0.8% agarose electrophoresis gel in 0.5% TAE (Tris-acetate-EDTA) buffer and visualized under ultraviolet light after ethidium bromide staining. Before Sanger sequencing, PCR products were purified using a Wizard® SV gel and PCR clean-up system kit (Promega, ref A9281). Sequences were cleaned by manually trimming low quality base pairs from the 3’ and 5’ ends using Geneious Prime (version 2.1) (https://www.geneious.com). We ensured the identity of the four *O.tauri* strains (RCC4221, RCC1116, RCC1108, RCC1123) by amplification of variable regions of chromosome 20 by PCR consisting of F36 (5′-TCG ACG CAC CTT TTC CCC-3′) and R641 (5′-CGA CGA CGA CGC TCG AAC-3′) primers.

### Concentration and purification of viral particles

A total of two liters of viral lysate from each strain was produced and filtered through a 0.2-µm PES membrane filter (SARSTEDT, ref 83.1826) to remove cellular debris and bacteria. Viral particles filtered were concentrated 40-fold by tangential flow filtration using a 100 kDa MWCO PES membrane Vivaflow200 unit (Sartorius, Germany, ref VF20P4). We concentrated the filtered lysate a second time with a 50,000 MWCO PES membrane Vivaspin20 ultracentrifugation unit (Sartorius, Germany, ref VS2031). Viral particles were further purified in an iodixanol (Optiprep™, Sigma-Aldrich, ref D1556) density gradient by ultracentrifugation rotor (SW41 Ti rotor, Beckman Coulter) at 175,117 × *g* for 5 h at 4°C. For each virus, a single visible band in the 30% fraction, corresponding to the purified concentrated viral particles, was collected and pelleted by ultracentrifugation at 245,000 *g* for 2.5 h at 4°C (SW55Ti rotor, Beckman Coulter). The virus pellets were resuspended in a sterile SM buffer (100 mM NaCl, 8mM MgSO4-7H2O, 50mM Tris pH 7.5) and stored at 4°C.

### Extraction and sequencing of viral DNA

DNA was extracted from the purified viruses with the CTAB protocol, as previously described (Winnepenninckx *et al*., 1993) with the following modifications. After CTAB buffer incubation (65°C for 30 min), an equal volume of phenol:chloroform:isoamyl alcohol (25:24:1) was added, and the samples were centrifuged (15,000 × *g* for 10 min at 4°C) . The aqueous phase was recovered, mixed with an equal volume of chloroform:isoamyl alcohol (24:1) and centrifuged (15,000 × *g* for 10 min at 4°C). 2 µL of RNAse A (1 M) (DNase free) was added to the aqueous phase for 30 min at 37°C, and then the enzymes were inactivated by heating for 10 min at 60°C. One tenth volume of ammonium acetate (3 M, pH 5.3) was added in order to favor DNA precipitation. Nucleic acids were precipitated using 2.5 volumes of absolute ethanol, gently homogenized and transferred at -80°C overnight. The precipitated DNA was centrifuged at 15,000 × *g* for 30 min at 15°C and the pellet washed in 70% ethanol. Finally, the samples were centrifuged (15,000 × *g* for 10 min at 15°C) air-dried and DNA dissolved in 50 µL of DNAse/RNAse free water and stored at 4°C. DNA in 1 µL was quantified by fluorimetry (Quantus™) using QuantiFluor® dsDNA system kit (Promega, ref E2670).

Viral DNA (1 µg) was submitted to Illumina NEXTFLEX library preparation (average insert size 241 bp) and sequenced using Illumina NextSeq 550 technology in paired-end mode (2×150 bp) giving a yield of 120 million reads total. Reads were cleaned with Trimgalore software (v0.6.5) (Krueger 2012) (with options --paired --fastqc --illumina --quality 26 --stringency 3 --length 35 -- three_prime_clip_R1 10 --three_prime_clip_R2 10 --clip_R1 10 --clip_R2 10) and the read quality evaluated with FastQC (v0.11.7)(Andrews 2010) with default settings.

### De novo genome assembly

We found that optimal *de novo* assembly of viral genomes required read data down-sampling, as using the entire dataset with estimated coverage of 6,000x induced assembly fragmentation. Down- sampling was performed using the tool seqtk version 1.3 (Li 2012). A total of eight independent sets of randomly down-sampled reads (seed numbers 100) were obtained given a coverage of 5x, 10x, 50x, 75x, 100x, 500x, 1,000x and 2,500x for each virus genomic library. *De novo* assembly was performed using Spades (v3.15.1) (Bankevich et al. 2012) all sets of independently down-sampled reads, as well as for the total set of cleaned reads. In order to ensure optimal assembly, and since some samples were revealed to have two independent viruses, we further assessed two assembly options: ‘--isolate’ and ‘-- meta’. The 18 assemblies for each viral strain were compared and the choice of the ‘best’ assembly was based on optimizing for 3 quality criteria given by QUAST (v5.0.0) (Gurevich et al. 2013). Specifically, we sought to maximize the size of the longest contig and to reduce ambiguous nucleotide positions (lowest number of N’s) and the total number of contigs. To facilitate the comparative genomic analysis, genomes comprising several contigs were manually scaffolded using Geneious (v11.0.3+7)(https://www.geneious.com), to ensure that the order of genes was conserved relative to that of viral genomes that were assembled as a single contig. The reads were mapped to the final genome assembly with the Burrows-Wheeler Aligner (BWA v0.7.17) using maximum exact matches (BWA-MEM) and a minimum seed length of 20 while keeping the other default parameters to further assess a correct assembly (Li & Durbin 2009). Specifically, we verified re-mapped read depth was even along the length of the genome.

### Gene prediction and functional annotation

Gene prediction of OtV genomes was performed using Prokka (v1.14.5) specifying ‘virus’ as the taxonomic kingdom (Seemann 2014). First, protein-coding genes (CDS) were annotated using the databases implemented in Prokka with the following parameters: e-value, 1e-5; and genetic code, standard (--gcode 1). Secondly, tRNA were predicted using tRNAscan-SE with default parameters (v2.0.2)(Chan & Lowe 2019). Thirdly, additional functional annotation of all protein sequences was sought by BLASTp search against the NCBI nr database accepting an e-value of <1e-5 keeping only the best hit. Finally, taxonomic affiliation was associated for each protein accession using Entrez Direct (EDirect) (epost -db protein | esummary -db protein | xtract -pattern DocumentSummary -element Caption,TaxId)(Kans 2021).

Structural predictions of single protein sequences from OtV genomes hypervariable regions were conducted with AlphaFold3 on the server https://alphafoldserver.com/ (version AlphaFold-beta- 20231127) (Abramson et al. 2024) with automated seed and otherwise standard parameters. Per-atom confidence estimates are recorded as pLDDT in the b-factor column of CIF files, with regions of pLDDT>90 representing very high confidence and regions of pLDDT<50 representing very low confidence. Best ranking PDB model for each target were then submitted to Foldseek structural similarity search on the server https://search.foldseek.com/search through all available databases (AlphaFold/Proteome, AlphaFold/Swiss-Prot, AlphaFold/UniProt50, CATH50, GMGCL, MGnify- ESM30, PDB100) in 3Di/AA mode (van Kempen et al. 2024). Most significant explicit descriptions of each target function were chosen based on the coverage of alignment of target and query, the statistical e-value of alignment, and Foldseek self-consistency between databases.

### Comparative genomic analysis

The complete genomes of the previously available prasinoviruses were downloaded from NCBI (Supplementary Table 2). A total of thrity-six genomes (nineteen new sequences genomes and seventeen prasinovirus references) were used to define groups of orthologous genes (orthogroups), where we excluded prasinovirus reference genomes that were incomplete or too fragmented (i.e., BII- V1, OlV3, OlV4, OlV5 and OlV6). All translated CDS were included in the orthogroup analysis using Orthofinder with default settings (Emms & Kelly 2019). Each orthogroup was annotated *via* the EggNOG-mapper toolkit (v2.0.4-rf1) against EggNOG protein database (-m diamond) using setting parameters (Huerta-Cepas et al. 2017). The Euclidean distance between the viruses based on the presence/absence pattern of orthogroups was calculated using pvclust R package (v2.2-0) based on the “ward.D2” hierarchical clustering linkage method and approximately unbiased (AU) *p*-values are computed using 1,000 bootstrap replicates (default settings). Orthogroup distribution in each genome was represented using the pheatmap R package (v1.0.12). Pairwise whole-genome alignments of OtVs were carried out using pyGenomeViz v1.0.0 with default parameters (https://github.com/moshi4/pyGenomeViz) and pgv-pmauve, part of pyGenomeViz was used for visualization of genome alignment with ‘nucleotide method’ as setting parameters. Average Nucleotide Identity (ANI) between all orthologous genes shared between all pairs of OtV genomes was computed using FastANI (v.1.32) (Jain et al. 2018). Genomic maps of the hypervariable region between specialist and generalist viruses were generated using Easifig (v2.2.2) (Sullivan et al. 2011).

### K_a_/K_s_ ratio calculation

The ratio of non-synonymous (*K_a_*) to synonymous (*K_s_*) substitutions of the targeted coding sequences in the HVR was computed using KaKs_Calculator2.0 (D. Wang et al. 2010). For this purpose, the ratio was estimated across different alignments for genes with sequences more than 80% of nucleotide identity. The KaKs_Calculator2.0 was run on each nucleotide alignment group separately through different methods (option: -m MA, MYN, LPB, MS and NG). The value of the ratio was deemed reliable if the models compared gave the same score, and the method with the highest *p*-value was then selected.

### Phylogenetic analysis

Phylogenetic reconstruction based on full-length DNA polB protein and nineteen core proteins shared between *Prasinovirus* and *Chlorovirus* genomes was performed. Protein sequences were aligned using MAFFT (v.7.313) with the L-INS-i algorithm (Katoh et al. 2002). We trimmed each alignment removing positions containing more than 80% gaps with Goalign (v0.3.2) (Lemoine & Gascuel 2021). Phylogenetic trees of single-proteins and concatenated proteins (comprising 5,483 amino acid positions) were built using Maximum-Likelihood (ML) method with IQ-TREE version 2.0.6 (Minh et al. 2020). For mixture model analysis, the LG+F+R4, LG+R3 and LG best models were chosen (option -m MFP) according to the Bayesian information criterion (BIC) for concatenated proteins and full-length DNA polB protein, respectively (Kalyaanamoorthy et al. 2017). The branch support values were computed from 1,000 replicates for the Shimodaira-Hasegawa (SH)-like approximation likelihood ratio test (aLRT) (Guindon et al. 2010) and 1,000 ultrafast bootstrap approximation (UFBoot) (Hoang et al. 2018). The trees were visualized with Interactive Tree Of Life (iTOL) v6 (Letunic & Bork 2007).

## Acknowledgements

We would like to thank Morgan Gaïa; Vladimir Daric and Hélène Mayer of the BSBII, Banyuls Bioinformatic Service; genotoul bioinformatics platform Toulouse Occitanie (Bioinfo Genotoul, https://doi.org/10.15454/1.5572369328961167E12) for providing help and computing and storage resources; and Christophe Salmeron and David Pecqueur from the SU/CNRS BioPIC Imaging and Cytometry platform of Banyuls-sur-Mer Oceanological Observatory, whose French state funds are managed by the ANR within the Investments of the Future program under reference ANR-10-INBS-02.

## Data availability

The genome sequences and annotations are available on Genbank with the following accession numbers: McV20-T, PQ442261; OtV06-1, PQ442262; OtV06-12, PQ442263; OtV06-4, PQ442264; OtV09-556, PQ442265; OtV09-557, PQ442266; OtV09-559, PQ442267; OtV09-565, PQ442268; OtV09-570, PQ442269; OtV09-573, PQ442270; OtV09-578, PQ442271; OtV09-582, PQ442272; OtV09-585, PQ442273; OtV19-O, PQ442274; OtV19-P, PQ442275; OtV19-R, PQ442276; OtV19-T1, PQ442277; OtV19-T2, PQ442278; OtV-Sylt2-5, PQ442279.

The complete list of AlphaFold3 models is available for download: (https://dropsu.sorbonne-universite.fr/s/8RREeAoBoDastwp).

