## Supplementary Figures and Table legends for "Genomic analysis of *Ostreococcus tauri*-infecting viruses reveals a hypervariable region associated with host–virus interactions"

**Supplemental data**

Supplementary Table 1. Summary of marine eukaryotic phytoplankton challenged for isolation of viruses from environmental seawater. Orange boxes correspond to sampling water collected from the south Pacific Ocean and the blue box from the North Sea. Virus-permissive strains are indicated by a plus (+) and non-permissive strains indicated by minus (-) signs. *strain used to isolate new viruses in this study.** New phytoplankton strains isolated. Med. Sea: Mediterranean Sea.

| Phytoplankton strains challenged for isolation of viruses | | | | Environmental from which viruses isolated | | | | |
|---|---|---|---|---|---|---|---|---|
| Phylum | Class | Strain | Origin | O | P | R | T | Sylt2-5 |
| Chlorophyta | Trebouxiophyceae | *Picochlorum* sp. F4_5** | NA | - | - | - | - | - |
| | Mamiellophyceae | *Bathycoccus prasinos* RCC4222 | Med. Sea | + | + | + | + | - |
| | | *Ostreococcus tauri* RCC4221 | Med. Sea | +* | +* | +* | +* | +* |
| | | *Micromonas commoda* RCC827 | Pacific Ocean | + | + | + | + | - |
| | | *Mamiella* sp. RCC391 | Atlantic Ocean | - | - | - | - | - |
| | | *Mantoniella* sp. RCC417 | Atlantic Ocean | - | - | - | - | - |
| | | *Mantoniella* sp . RCC6849 | Med. Sea | - | - | - | - | - |
| | Pyramimonadophyceae | *Pyramimonas* sp. RCC6848 | Med. Sea | - | - | - | - | - |
| Haptophyta | Pavlovophyceae | *Pavlova* sp . A511** | NA | - | - | - | - | - |

Supplementary Table 2. Genomes characteristic of prasinoviruses used in the present study. Virus names in bold correspond to prasinoviruses sequenced and characterized in this study. Note that the *O. tauri* strain RCC4221 was cloned from RCC745.

Supplementary Table 3. Orthologous protein-coding genes in *Prasinovirus*. a) The orthogroups (OG) matrix shared between the prasinovirus genomes is shown in sheet 1 and b) the locus associated in sheet 2. c) The unique genes in the viral genomes are indicated in the sheet 3 with d) the locus tag associated in the sheet 4. Functional categories of orthogroups shared between the genomes. COG: Clusters of Orthologous Groups. e) Summary of the number of core and clade-specific orthogroups found among the prasinovirus genomes. "Core" refers to orthogroups that are common across a clade and present in 100% of genomes within an indicated group, while "clade-specific" corresponds to orthogroups only present in a specific clade of viruses and present at least in two genomes.

Supplementary Table 4. Pairwise alignment comparison of whole-genome Average Nucleotide Identity (ANI) of *Ostreococcus* tauri viruses.

Supplementary Table 5. Analysis of selection pressure across pairwise nucleotide alignments within the HVR. Group: different alignments of orthologs which each have ≥80% aa identity. Putative function: Predicted gene functions. Gene_locus sequence: Locus of gene sequences in the nucleotide alignment. Length: Length of the nucleotide alignment in base pairs (bp). Model: NG: Nei–Gojobori; MA: Modified Approximate; MS: Modified Smith; MYN: Modified Yang & Nielsen; LPB: Li, Pamilo, and Bianchi. Ka: non-synonymous site. Ks: synonymous site. Ka/Ks: the ratio of non-synonymous polymorphisms per non-synonymous site to synonymous polymorphisms per synonymous site. p-value: p-value computed by Fisher's exact test. Type of selective pressure: purifying (negative) or diversifying (positive) selection.

Supplementary Table 6. Annotation of predicted genes of the hypervariable region. a) using BLASTX against the NCBI nr database. The top three best hits with an e-value <10-5 were displayed. qstart: Query alignment starts; qend: Query alignment end; sstar: Subject alignment start; send: Subject alignment end; e-value: expect value; stitle: Subject title. b) using BLASTP against the Refseq, Interproscan databases and Conserved Domains Database (CDD). E-value <10-5. c) from Alphafold 3.

Supplementary Table 7. Matrix presence/absence of the nineteen virus genes used to build the phylogenetic tree. Bold virus name corresponds to virus characterized in this work

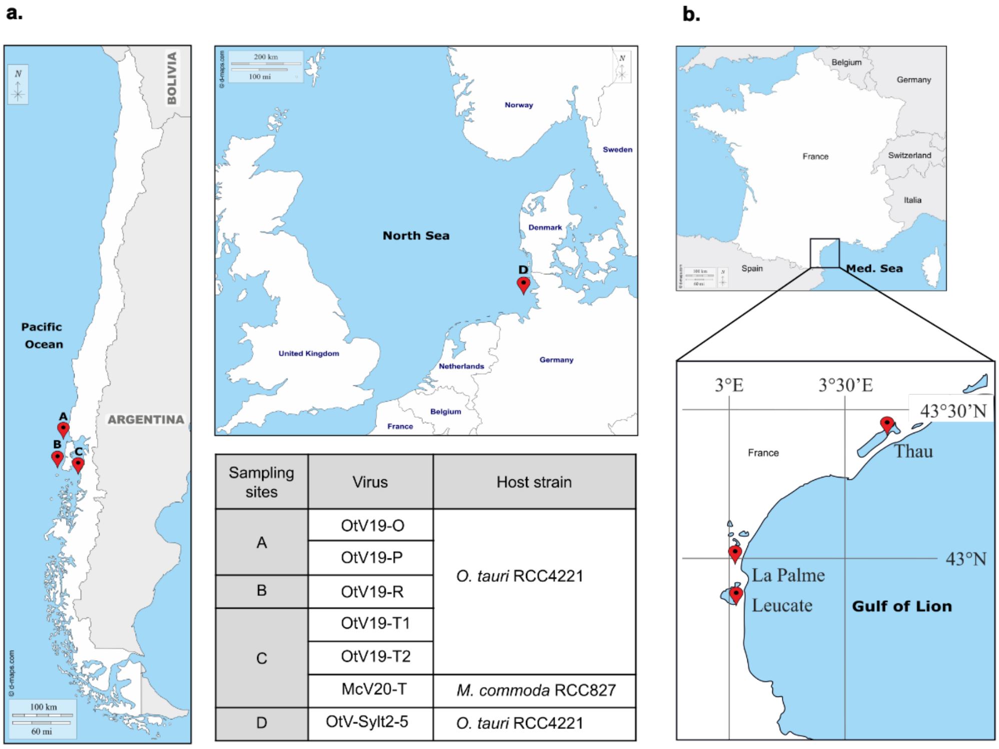


| Sampling sites | Virus | Host strain |
|---|---|---|
| A | OtV19-O | *O. tauri* RCC4221 |
| | OtV19-P | |
| B | OtV19-R | |
| C | OtV19-T1 | |
| | OtV19-T2 | |
| | McV20-T | *M. commoda* RCC827 |
| D | OtV-Sylt2-5 | *O. tauri* RCC4221 |

Supplementary Figure 1. Seawater sampling sites. a) Seawater was collected at three different sites (A, B, and C) on the coast of Chile in the Pacific Ocean and one site (D) in the North Sea along the German coast plotted in red on the map. Six new viruses were isolated from the *Ostreococcus tauri* RCC4221 strain, while one virus was isolated from the *M.commoda* RCC827 strain. b) Sampling sites located in the Mediterranean Sea and lagoons in southern France where twelve OtVs had been isolated in a previous study (Clerissi *et al.*, 2012). The geographical coordinates for each site and each virus are found in supplemental data (Table S1).

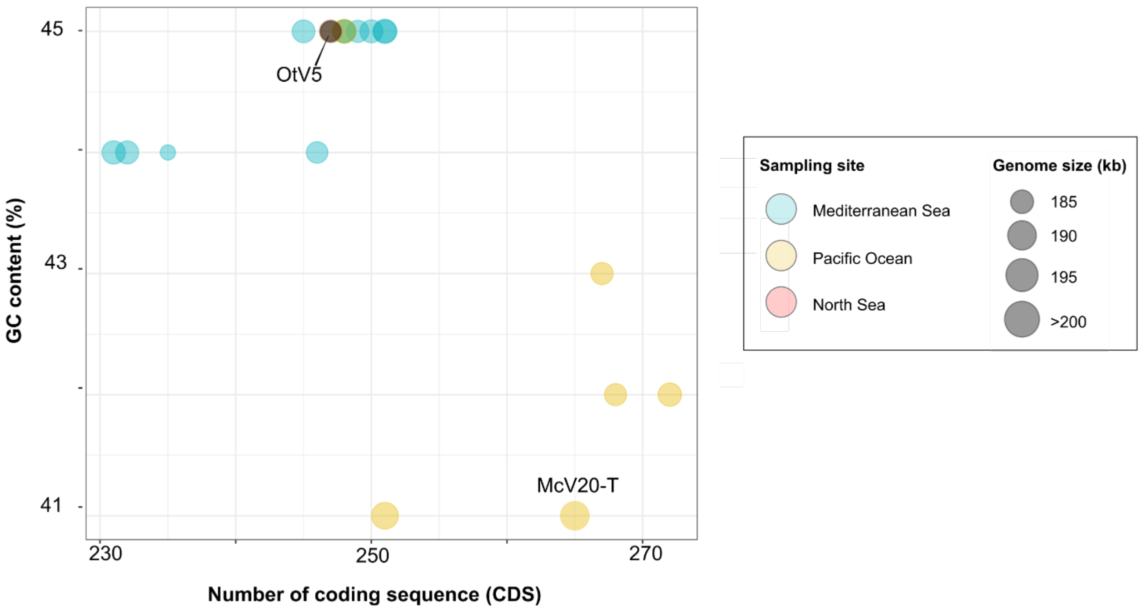


Supplementary Figure 2. Genomic characteristics of the new *Prasinovirus* sequenced. The x-axis indicates the number of coding sequences (CDSs) and the y-axis represents the GC content of the sequenced genomes. The size of the circles represents the genome size (in kb). Colors represent their geographical origin: Blue; Mediterranean Sea, Yellow; Pacific Ocean, Red; North Sea. The OtV5 virus was added as a reference. Raw data are shown in supplemental data (Table S2).

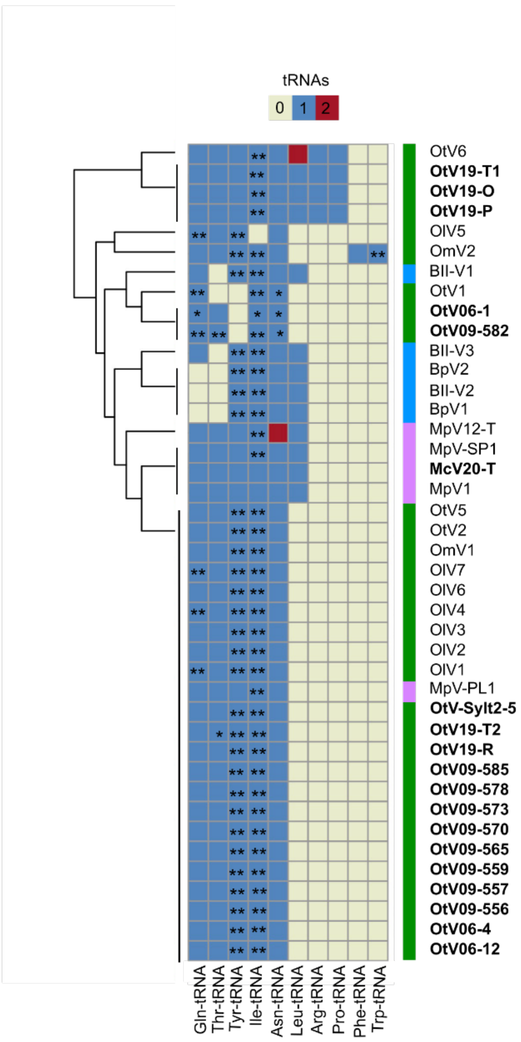


Supplementary Figure 3. Clustering based on tRNA patterns in the prasinovirus genomes. The x-axis indicates tRNAs and the y-axis the virus genomes containing tRNAs. Bolded names show the new viruses sequenced in this study. The host genus is reported by the strip colored: green; *Ostreococcus*, purple; *Micromonas* and blue; *Bathycoccus*. The number of tRNAs in each genome is indicated by colors: White; zero (0), Blue; one (1), Red; two (2). *:Predicted as tRNA pseudogenes; **:tRNA with introns.

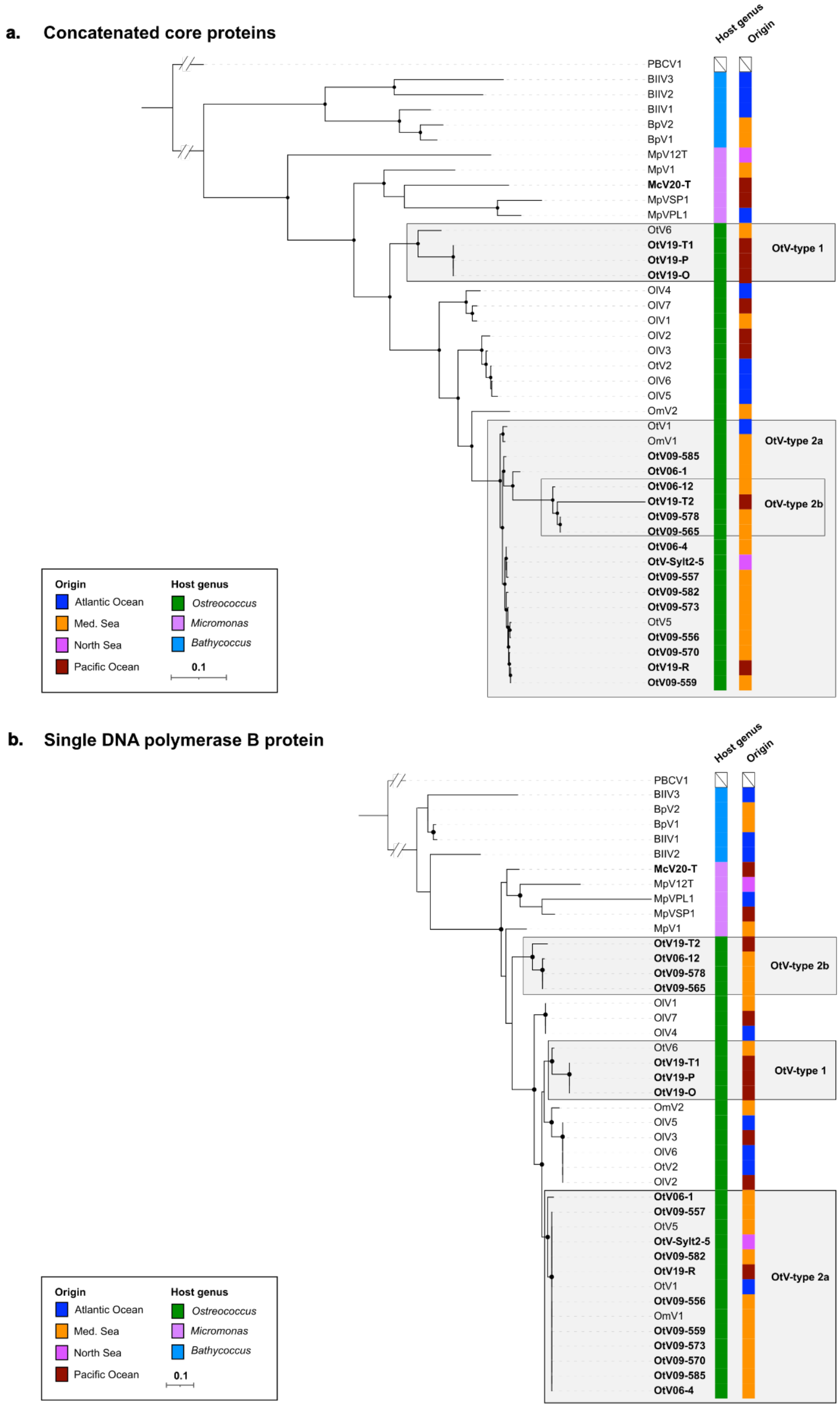


Supplementary Figure 4. Phylogenetic relationship between *Prasinovirus*. a) Maximum-likelihood (ML) phylogenetic reconstruction of nineteen core proteins shared among the viruses (5,529 amino-acid (aa) sites) infecting the genera *Bathycoccus*, *Micromonas* and *Ostreococcus* and *Chlorovirus* (PBCV1), which was used as an outgroup with the connecting branch truncated for display. The LG+F+R4 model was used based on the Bayesian Information Criterion (BIC). The support nodes were calculated by the SH-like aLRT (1,000 replicates) and UFBoot (1,000 replicates). Presence/absence matrix of core proteins for building the tree is shown in Table S7. b) Maximum-likelihood (ML) phylogenetic reconstruction of full-length DNA polymerase B protein (135 aa sites). The Chlorovirus (PBCV-1) was used as an outgroup with the branch connecting truncated for display. The LG+F+R3 model was used based on the Bayesian Information Criterion (BIC). The support nodes were calculated by the SH-like aLRT (1,000 replicates) and UFBoot (1,000 replicates). Nodes with bootstrap support over 80% are indicated by black dots. The scale bar represents the average number of substitutions per site. The host genus and the geographical origin are indicated by color stripes. Viruses belonging to each OtV-type (OtV-type 1, OtV-type 2a and OtV-type 2b) are delimited by gray boxes.

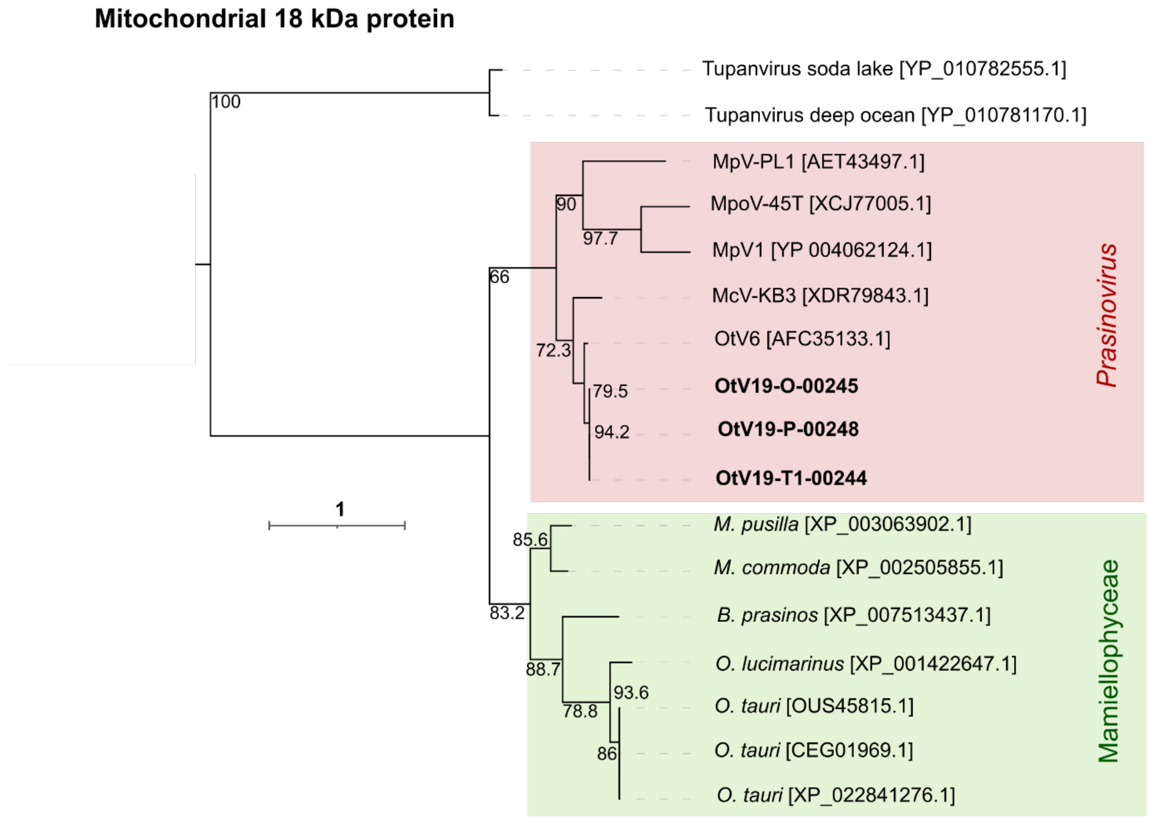


Supplementary Figure 6. The Maximum-likelihood (ML) phylogenetic tree of mitochondrial 18kDa protein was built using Q.pfam+G4 best-fit model based on the Bayesian Information Criterion (BIC). The support nodes were calculated by the SH-like aLRT (1,000 replicates) and UFBoot (1,000 replicates). The scale bar represents the average number of substitutions per site. New viruses isolated in this study are bolded in black. The prasinovirus clade is highlighted in red and the Mamiellophyceae host clade in green. The Tupanviruses branch was used to root the three.

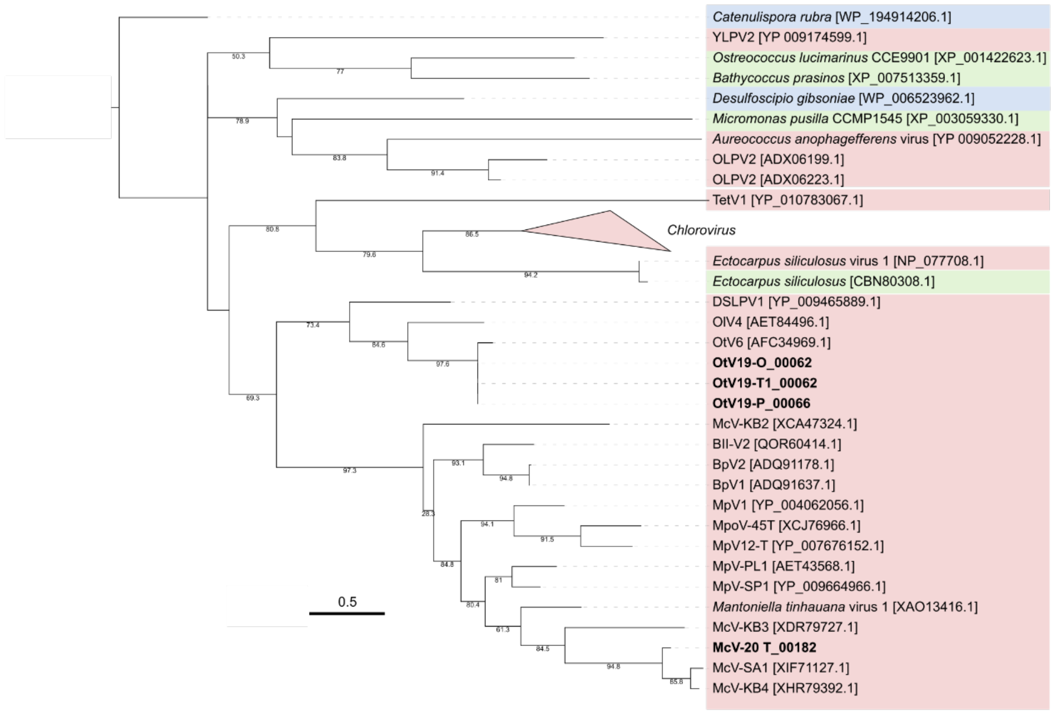


Supplementary Figure 7. Maximum-likelihood (ML) phylogenetic tree of the potassium channel protein was built with the best mtZOA+G4 model based on the Bayesian Information Criterion (BIC). Nodes with bootstrap support are indicated under the branches. The scale bar represents the average number of substitutions per site. Viruses isolated in this study are bolded in black. Virus branches are indicated in red, eukaryote branches in green and bacteria branches are in blue.

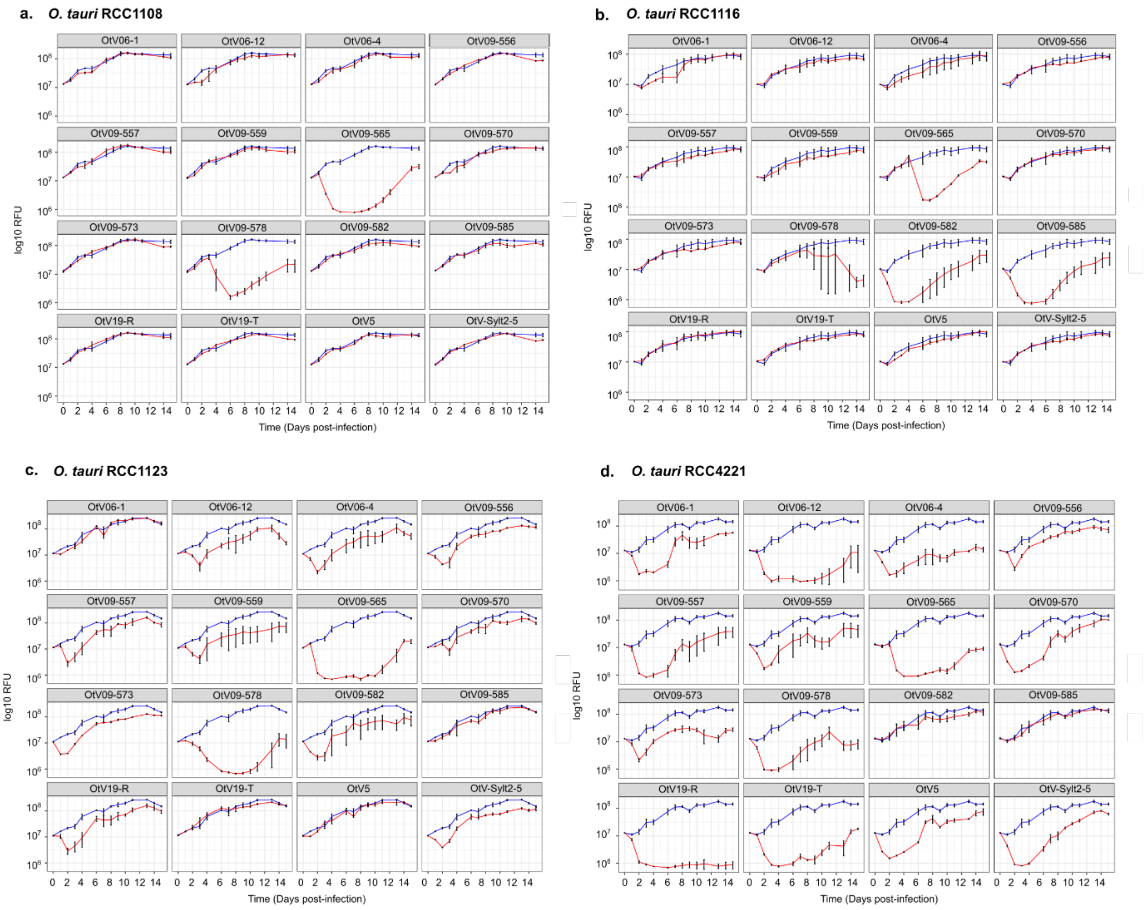


Supplementary Figure 8. Infection dynamics between *Ostreococcus tauri* strains and their viruses. *Ostreococcus tauri* a) RCC1108, b) RCC1116, c) RCC1123, d) RCC4221 strains. The log10-transformed chlorophyll fluorescence (Relative fluorescence unit — RFU) was measured over the time (days post-infection) in mock-inoculated cultures (control) (blue curve) and infected cultures (red curve). Each panel corresponds to different viruses (OtV06-1, OtV06-12, OtV06-4, OtV09-556, OtV09-557, OtV09-559, OtV09-565, OtV09-570, OtV09-573, OtV09-578, OtV09-582, OtV09-585, OtV19-R, OtV19-T, OtV5, OtV-Sylt2-5) used for the infection assay.

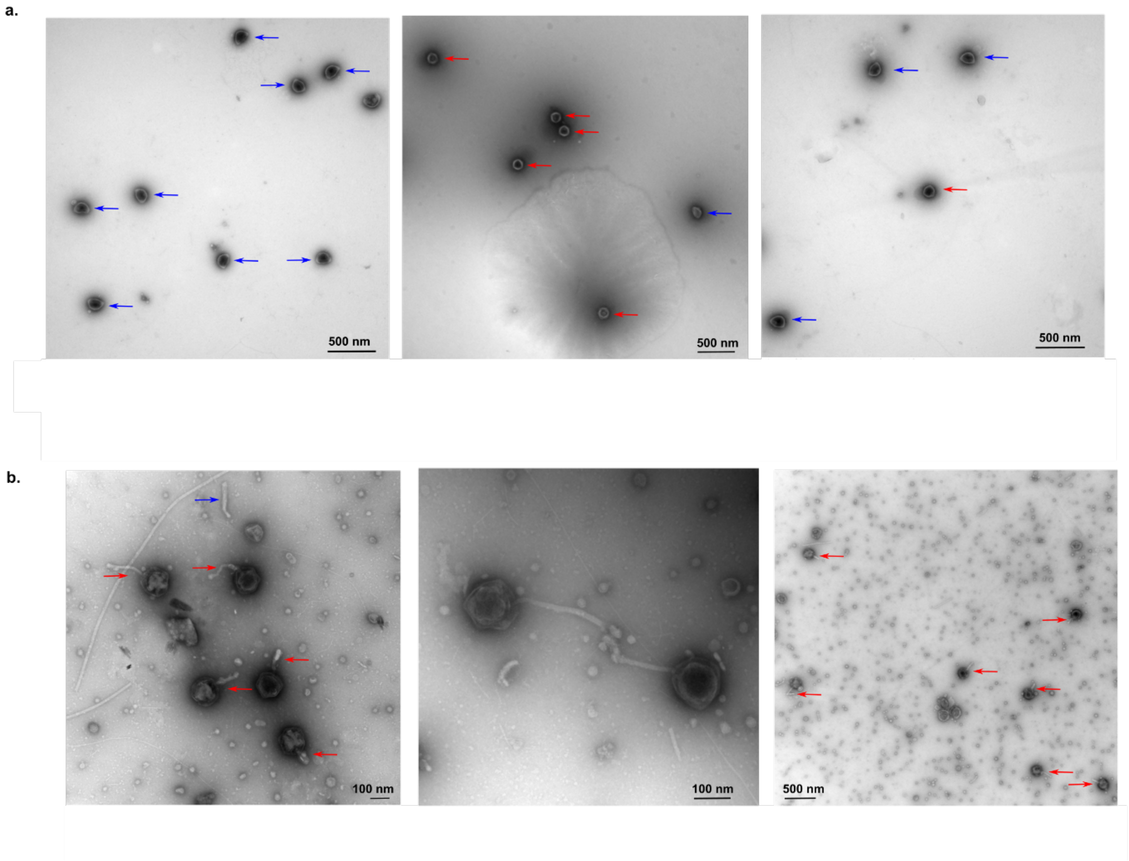


Supplementary Figure 9. Transmission electron micrograph of OtV like-particles. a) OtV19-T lysate consists of two distinct morphotypes of virions; non-enveloped icosahedral (red arrow) and "squashed" particles (blue arrow). b) OtV09-565 lysate showing the presence of viral particles with tubular structures like tails (red arrow) associated with the icosahedral capsid. The virions were negatively stained with uranyl acetate.

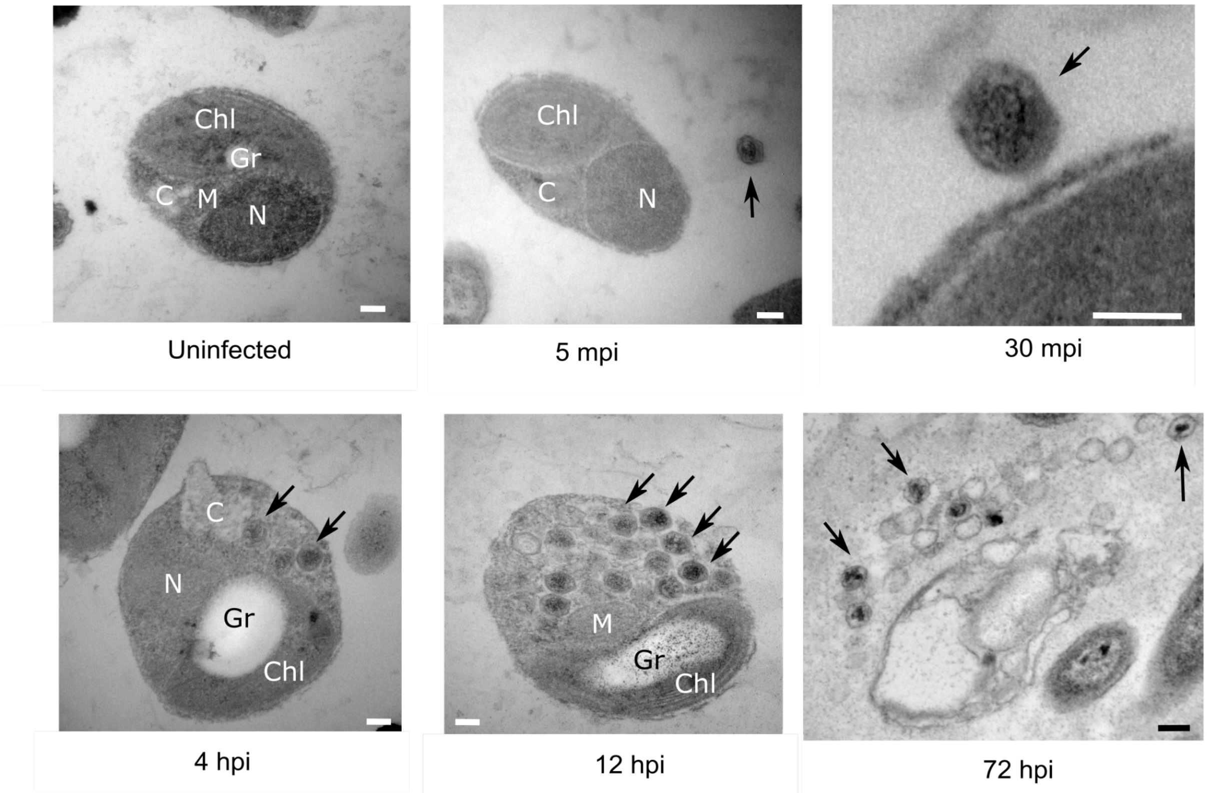


Supplementary Figure 10. Electron microscopy of thin sections of infected *O. tauri* RCC4221 cells with OtV09-565 virus over time; 5 mpi, 30 mpi (minute post-infection), 4 hpi, 12 hpi and 72 hpi (hour post-infection). The uninfected control culture was sampled at 30 mpi. Scale bar — 100 nm. Black arrows indicate viral particles. Chl - Chloroplast, N - Nucleus, M - Mitochondrion, C - Cytoplasm, Gr - Starch grain. Observations at 30 min show a filled virus particle attached to the plasma membrane of the host cell.

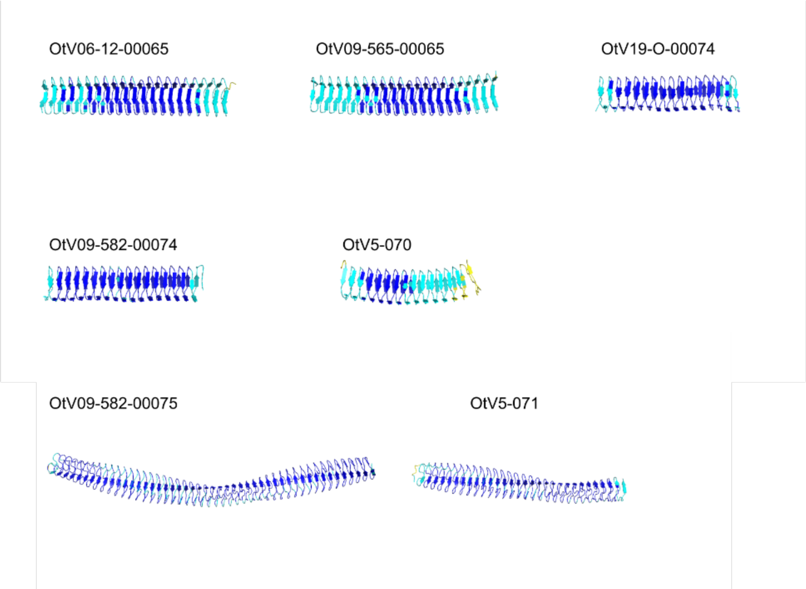


Supplementary Figure 11. Close-up view on β-folded segments of the three-dimensional (3D) of hypervariable sequences annotated as cell wall surface anchor family protein (FhaB) in the OtV06-12, OtV09-565, OtV5, OtV09-582, and OtV19-O genomes used as references. The AlphaFold models are colored according to prediction score; higher confidence residues in dark blue and cyan and lower confidence in yellow and orange.
