## Supplementary Figure 1 for "Genomic analysis of *Ostreococcus tauri*-infecting viruses reveals a hypervariable region associated with host–virus interactions"

**a.**

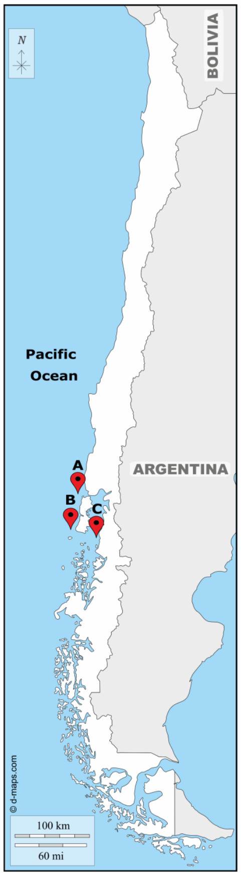


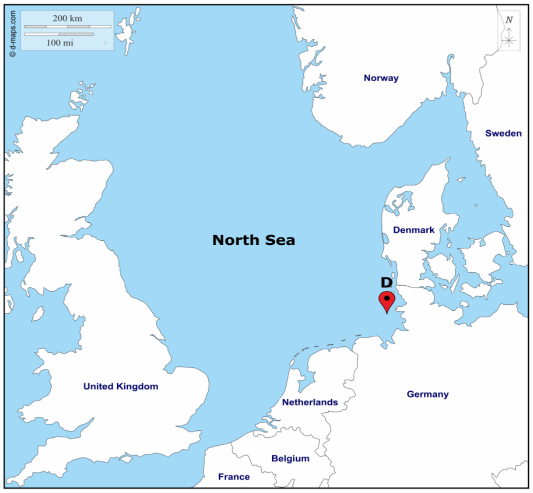


| Sampling sites | Virus | Host strain |
|---|---|---|
| A | OtV19-O | *O. tauri* RCC4221 |
| | OtV19-P | |
| B | OtV19-R | |
| C | OtV19-T1 | |
| | OtV19-T2 | |
| | McV20-T | *M. commoda* RCC827 |
| D | OtV-Sylt2-5 | *O. tauri* RCC4221 |

**b.**

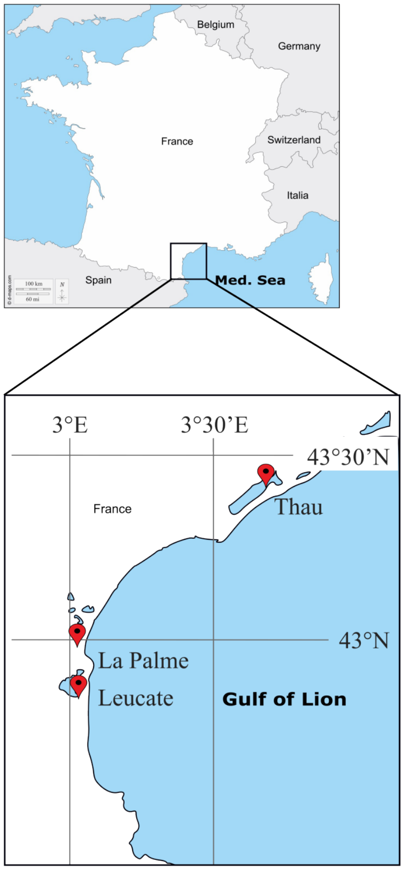
