## Supplementary figures and images for "Genomic analysis of *Ostreococcus tauri*-infecting viruses reveals a hypervariable region associated with host–virus interactions"

### Supplementary Figure 2

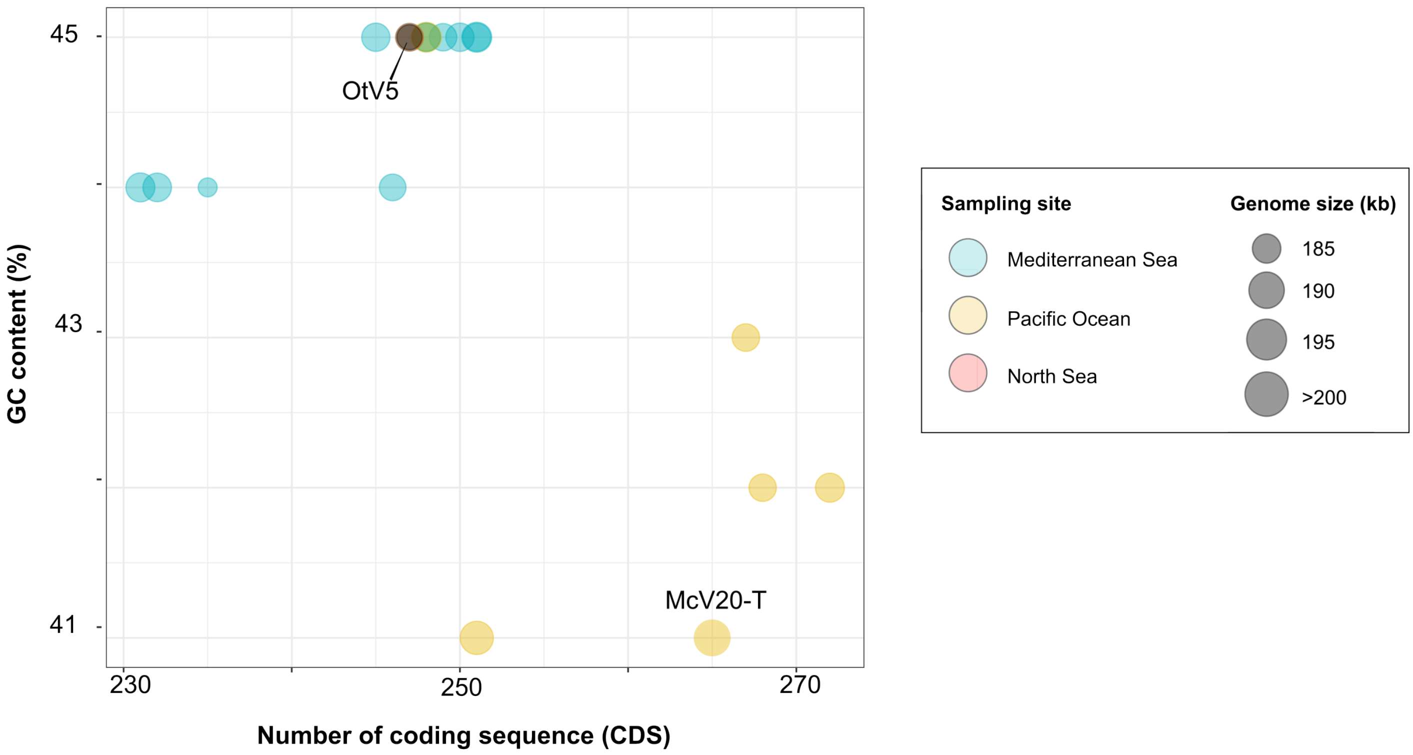

### Supplementary Figure 3

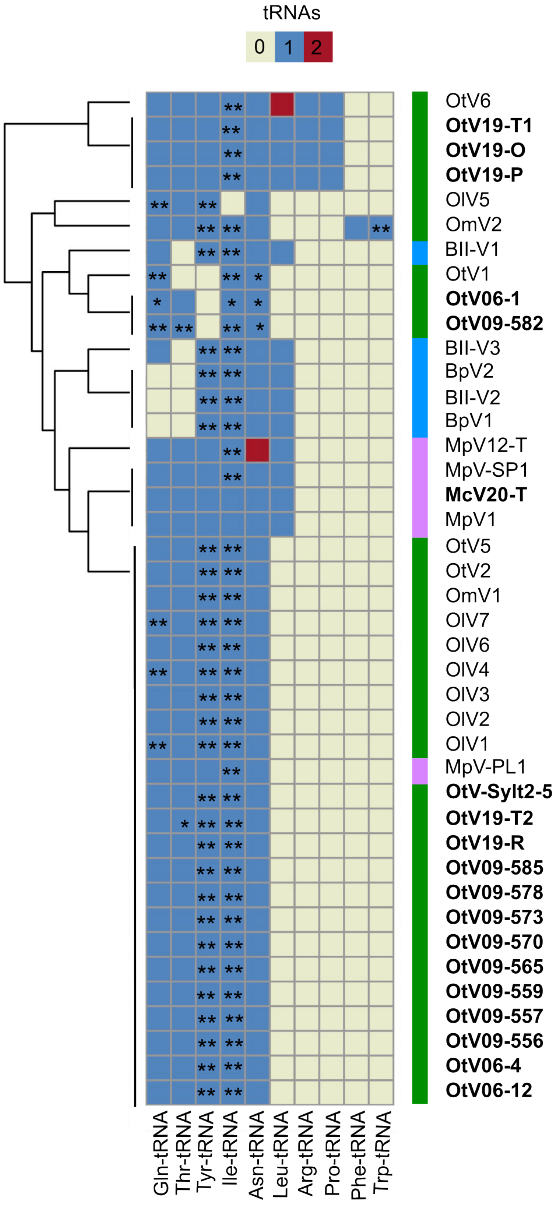

### Supplementary Figure 4

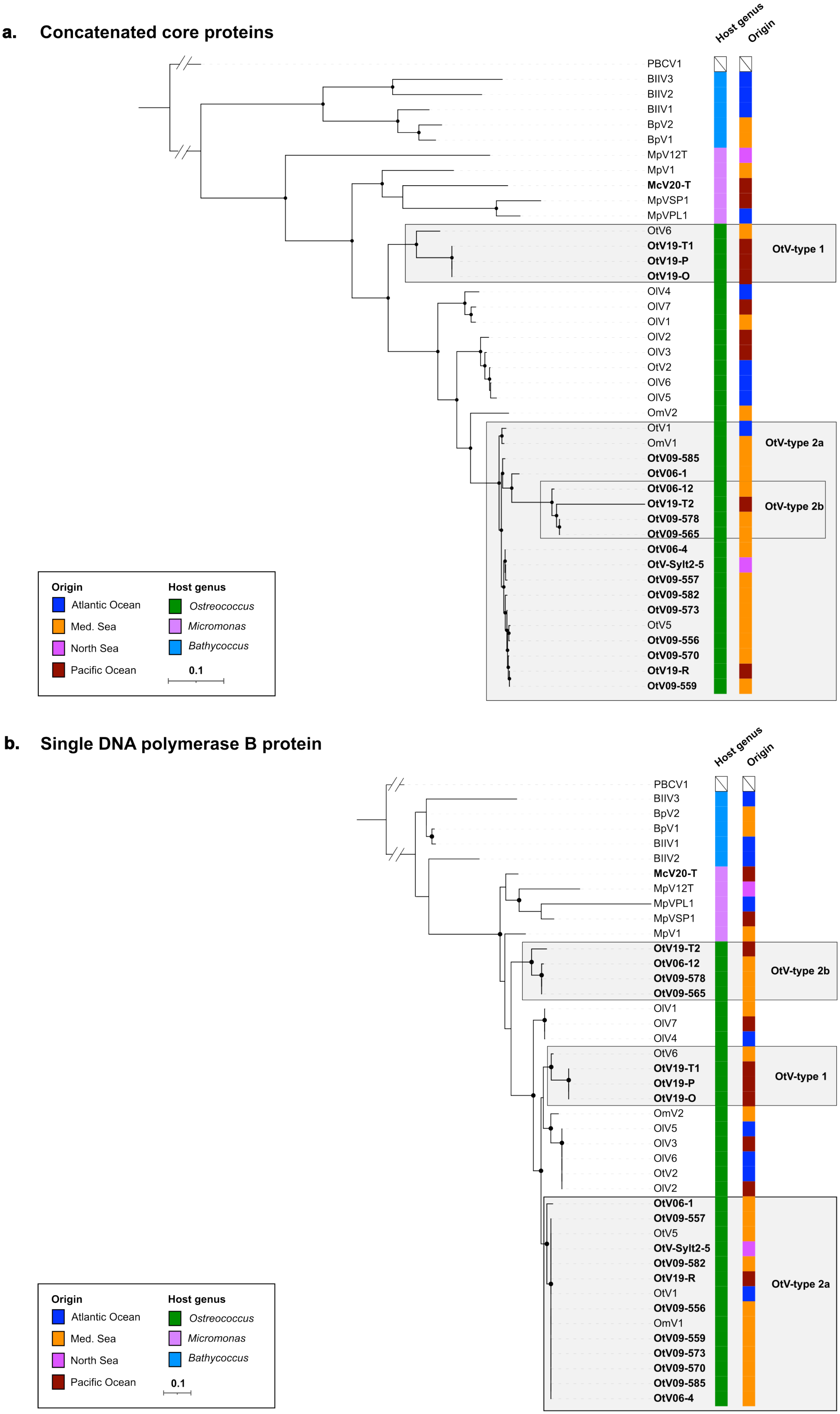

### Supplementary Figure 5

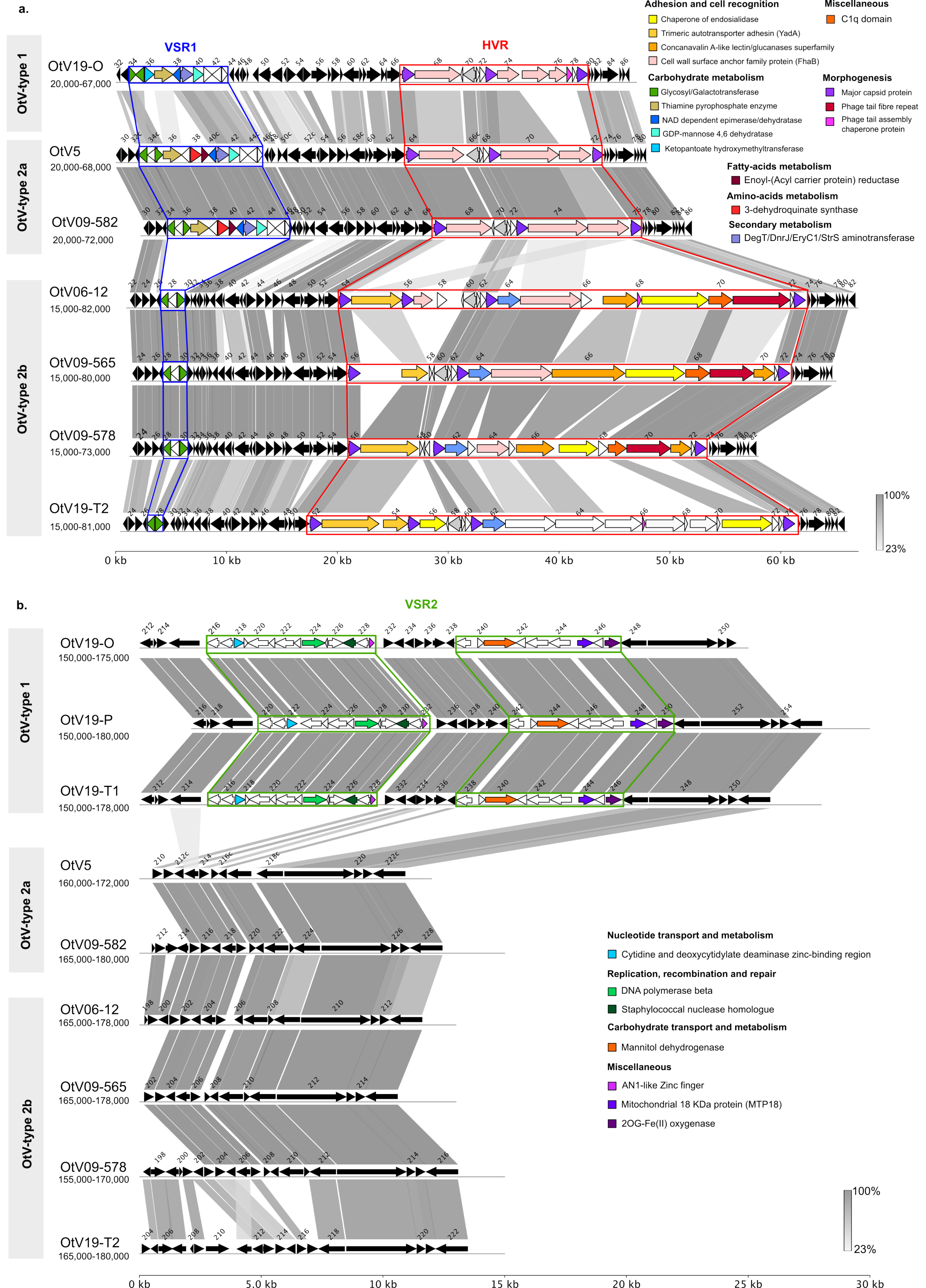

### Supplementary Figure 6

#### Mitochondrial 18 kDa protein

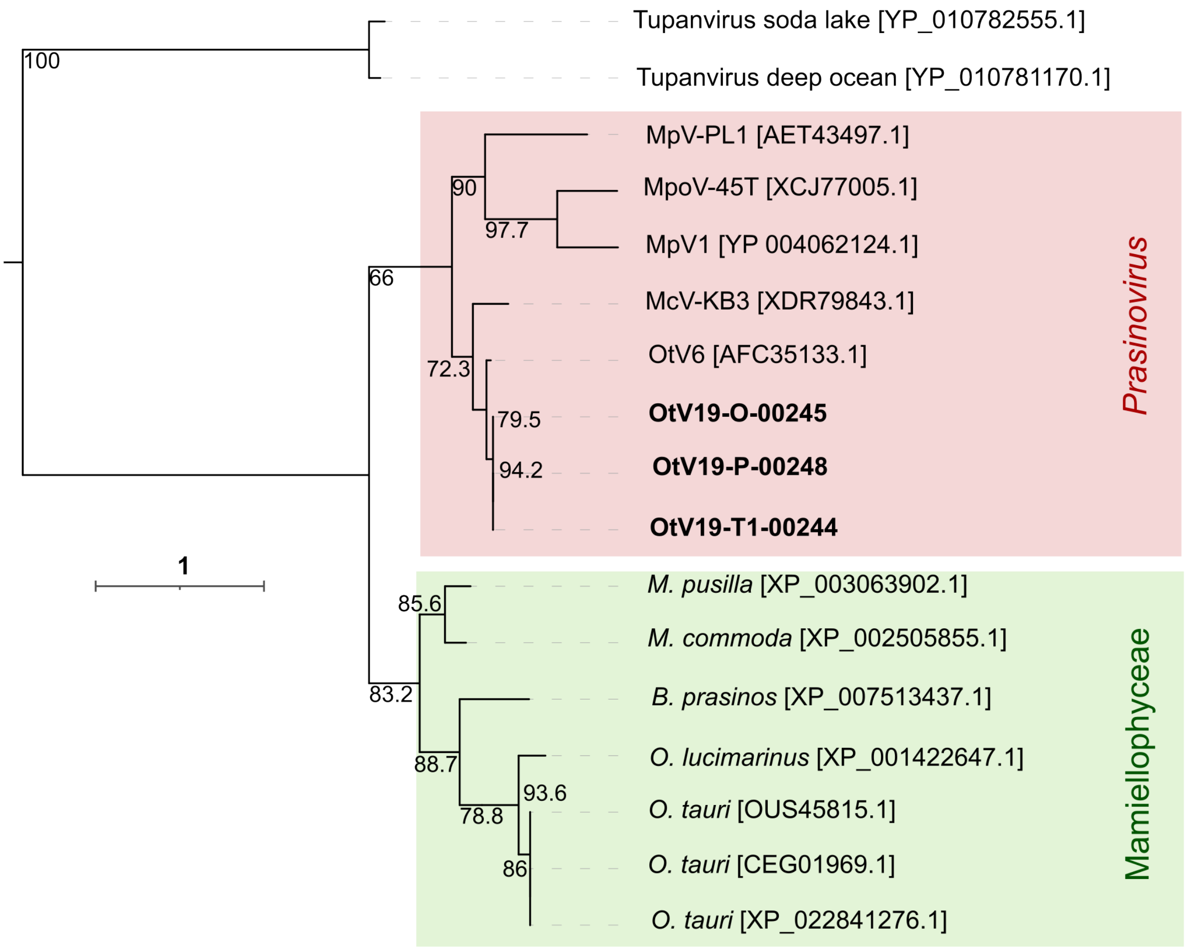

### Supplementary Figure 7

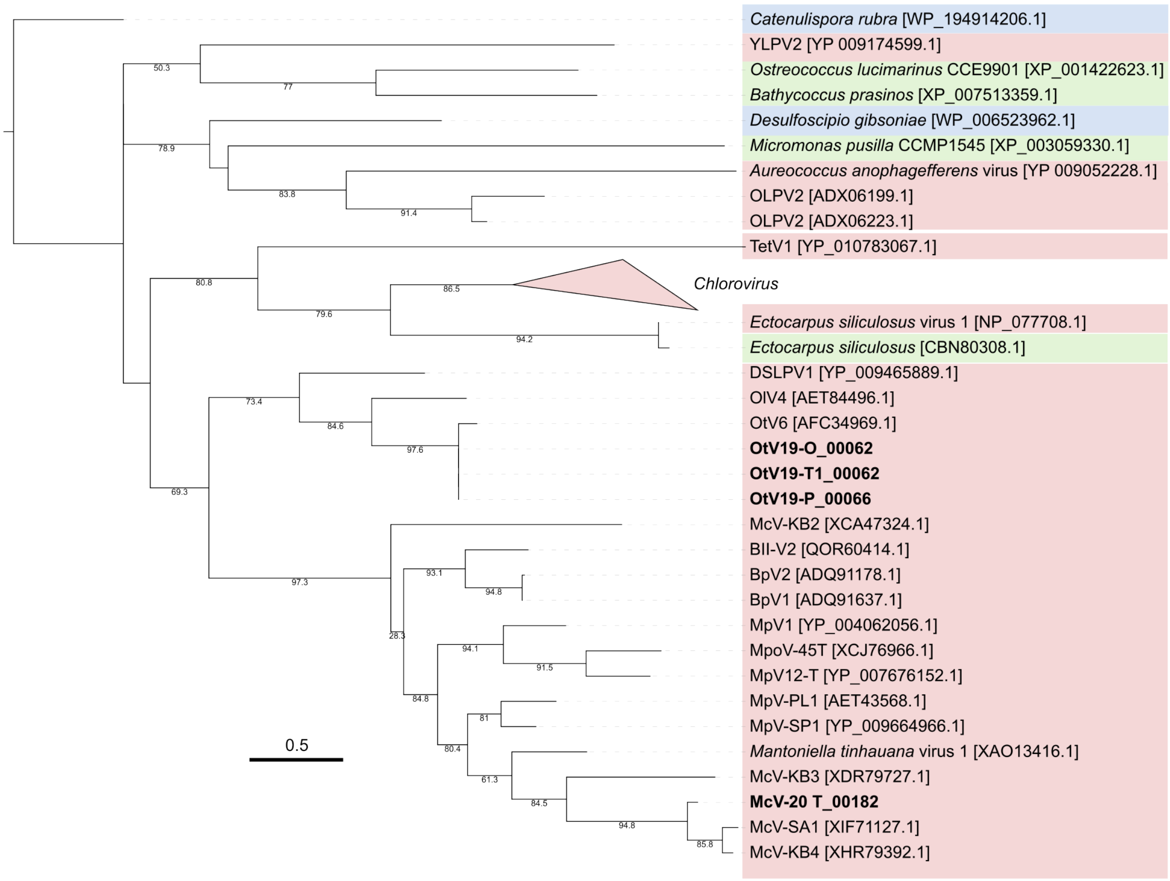

### Supplementary Figure 8

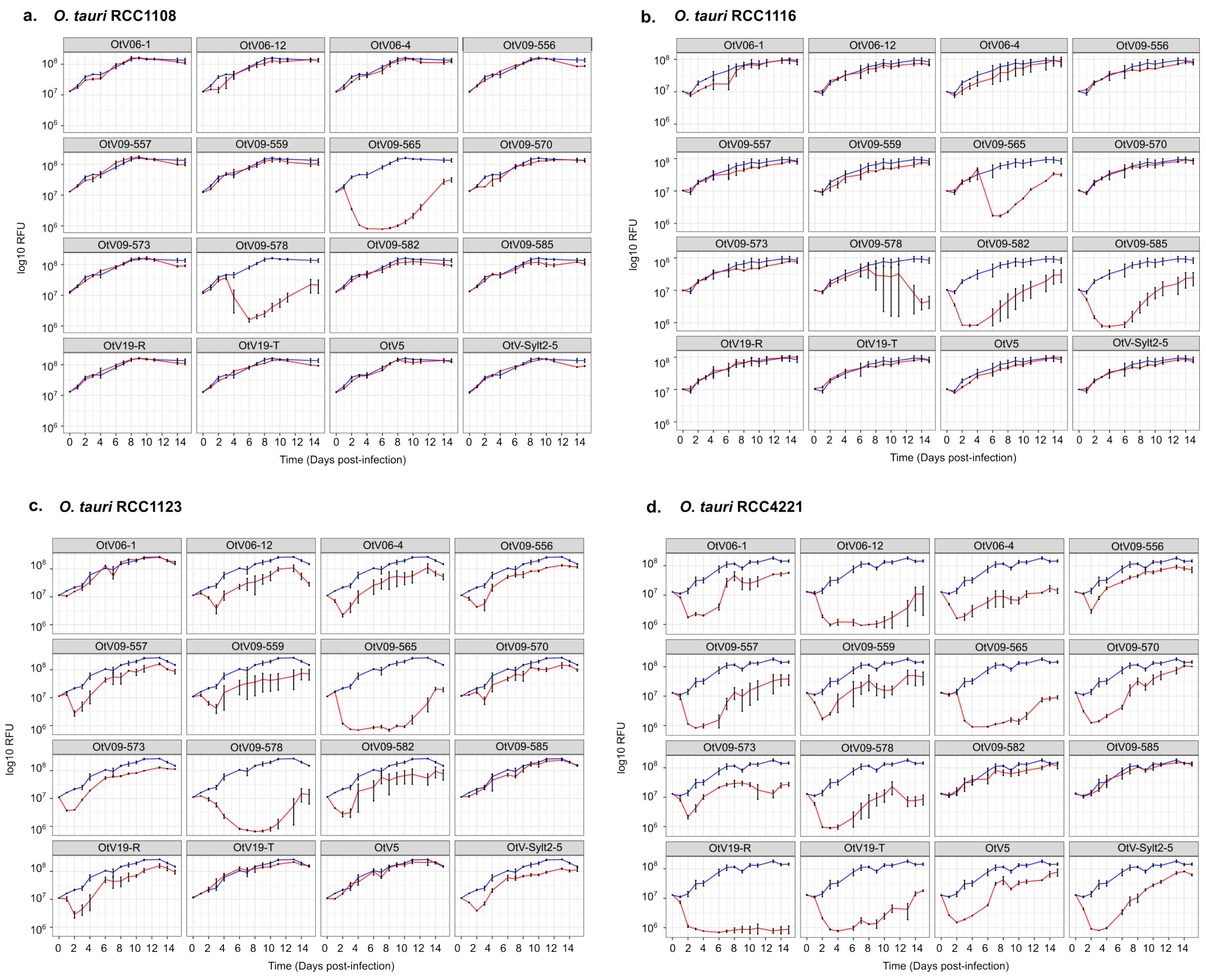

### Supplementary Figure 9

a.

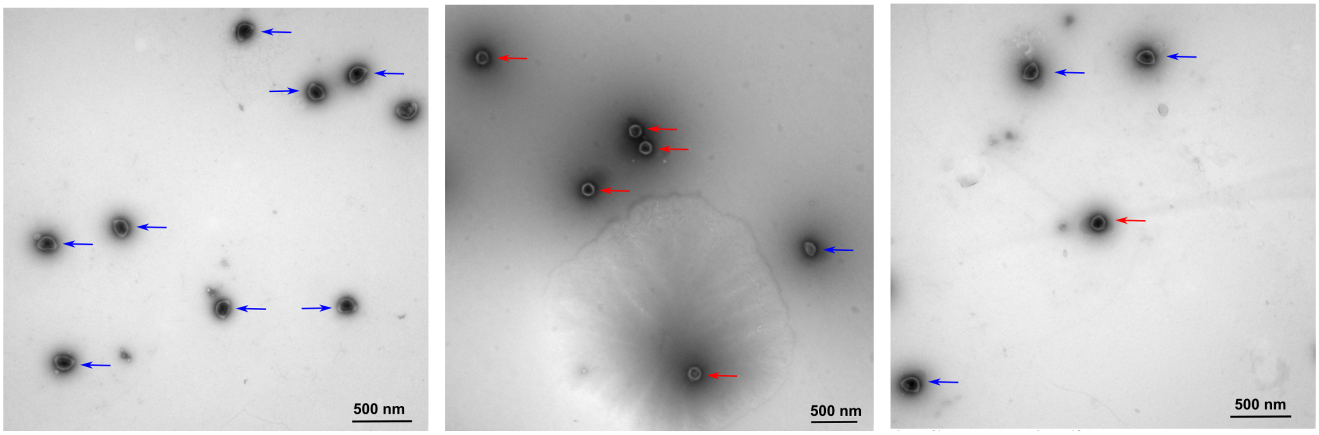


b.

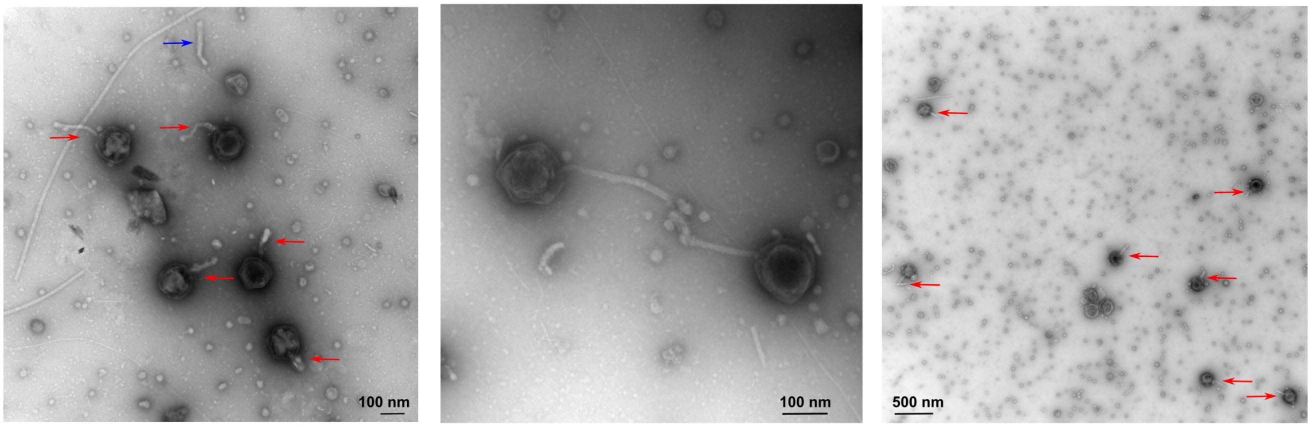

### Supplementary Figure 10

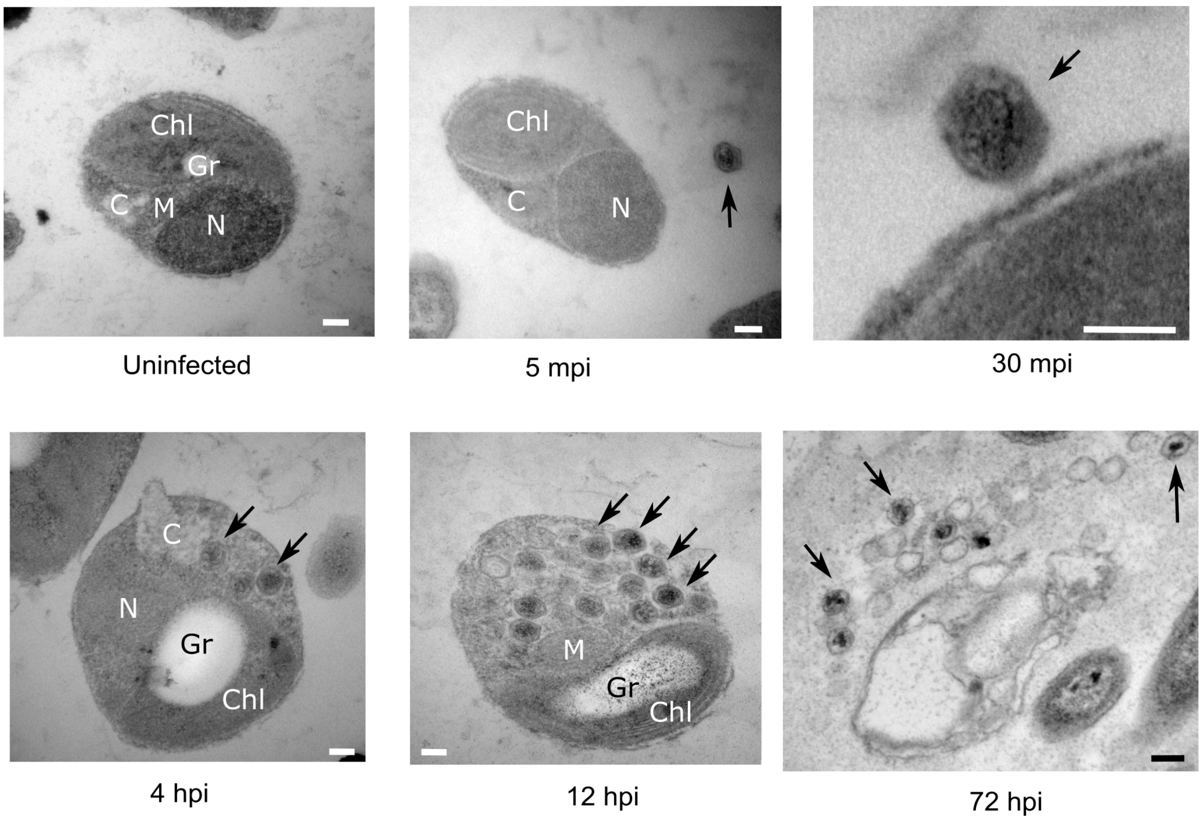

### Supplementary Figure 11

OtV06-12-00065

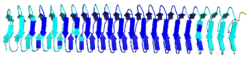

OtV09-565-00065

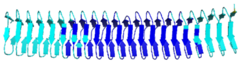

OtV19-O-00074

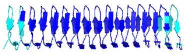

OtV09-582-00074

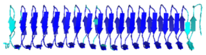

OtV5-070

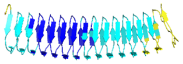

OtV09-582-00075

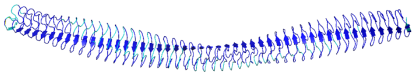

OtV5-071

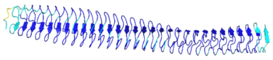
